# Implantable CMOS Deep-Brain Fluorescence Imager with Single-Neuron Resolution

**DOI:** 10.1101/2025.06.03.657675

**Authors:** Sinan Yilmaz, Jaebin Choi, Ilke Uguz, Jongwoon Kim, Alejandro Akrouh, Adriaan J. Taal, Victoria Andino-Pavlovsky, Heyu Yin, Jason D. Fabbri, Laurent Moreaux, Michael L. Roukes, Kenneth L. Shepard

## Abstract

Despite the advantages of optical imaging over electrophysiology, such as cell-type specificity, its application has been limited to the investigation of shallow brain regions (< 2 mm) because of the light scattering property of brain tissue. Passive optical conduits such as graded-index lenses and waveguides have permitted access to deeper locales but with restricted resolution and field of view, while creating massive lesions along the inserted path, with little pathway to improvement in the technology. As an alternative, we present the Acus device, an active implantable complementary metal-oxide-semiconductor (CMOS) neural imager with a 512-pixel silicon image sensor post- processed into a 4.1-mm-long, 120-μm-wide shank with a collinear fiber for illumination, which is able to record transient fluorescent signals in deep brain regions at 400 frames/sec. Acus can achieve single-neuron resolution in functional imaging of GCaMP6s-expressing neurons at a frame rate of 400 frames/sec.

## INTRODUCTION

Deciphering neural activity on a large scale within the brain is crucial to the discovery of the neural mechanisms underlying complex behavior and neurological disorders.^1,2^ Current methods for functional recording of cortical activity are dominated by electrophysiology (ephys). Integrated circuits based on complementary metal-oxide-semiconductor (CMOS) technology have provided a successful path for microfabrication of implanted electrode arrays, enabling both the scaling of the number of recording sites per shank and the precise dimensional control of spacing and configuration of recording sites.^3^ The most successful example of an implantable ephys array is the Neuropixel shank design.^4,5^

Despite this progress, electrical recording approaches have some notable weaknesses^6^; in particular, the circuit complexity of ephys recording channels exceeds that required of optical detectors, allowing the latter to be integrated at high densities. As a result, for the same displaced volume, more neurons can be recorded with an optical shank than an ephys one. Optical techniques can also interact with neurons at greater distances than ephys interfaces allowing large effective tissue volumes to be interrogated. In addition, precise identification and control with cell-type specificity are lacking in ephys approaches. Using electrodes to identify different types of cells in local brain circuits, such as pyramidal cells and interneurons, depends on analyzing spike waveforms and other indirect measures that are relatively unreliable.^7–11^ Using electrodes to identify neurons that form large-scale networks by sending projections to other brain regions depends on low-yield collision tests.^12,13^ These limitations mean that light and microscopy have real advantages in recording neural activity since cell types can be discerned with labelling and activity can be detected at distances further from the detectors.

Traditionally, optical imaging of neurons has been accomplished using microscopy techniques like multi-photon microscopy which has demonstrated excellent spatial resolution with head-fixed animal models.^14^ To enable long-term experiments crucial for behavioral tests, lens-based microscopes have been miniaturized into a fully implantable form-factor.^15^ However, one main challenge with existing optical techniques is their probing depth. Light scattering and absorption in tissue fundamentally limit the depth at which fluorescence microscopes can detect labeled neurons with the depth of light penetration typically restricted to less than 2 millimeters, even when using infrared wavelengths.^16,17^ Existing approaches for enhancing the depth capabilities of microscopy techniques, including fiber photometry^18–20^, micro-endoscopes^21,22^, or the integration of graded-index (GRIN) lenses^23–25^ provide restricted fields-of-view (FoV) with large cross sections for implantation damage.

An alternative to these approaches to achieve imaging at depth is to implant an optoelectronic device to record fluorescent signals from neurons in deep volumes directly. We have previously highlighted several critical challenges that must be addressed to develop a system capable of achieving single-cell resolution at depth,^6^ which include the effective rejection of background excitation light and the minimization of tissue damage during device insertion. The integration of volumetrically efficient, high-performance spectral and temporal filters is essential to address the first challenge. To address the second challenge, nanofabrication of the imager into a form factor that minimizes tissue disruption is necessary for successful device implementation. Additionally, integrating a method for light delivery significantly impacts both of these challenges. In this work, we address all of these issues through a combination of post-processing of the CMOS die, heterogeneous integration, and advanced optical packaging.

To accomplish this, we present in this work the elements of a lensless cellular-level imaging system, including electro-optical detectors to measure fluorescent signals, spectro-temporal filters and electronic circuits for control and data conversion, all fabricated on a 4.1-mm-long shank, which we call Acus, with a cross-sectional dimension of only 70 μm × 120 μm. Through a custom blind source separation (BSS) algorithm, we demonstrate the ability to localize fluorescently labeled sources in 3D space while having the ability to achieve imaging frame rates up to 400 frames/sec, consistent with the temporal requirement of recently developed genetically encoded voltage indicators (GEVIs).^26,27^ In our *in vivo* studies, we were able to image population dynamics at depth in densely labelled transgenic GCaMP6f-expressing mice, which demonstrates Acus’ capability to capture rapid neural dynamics. We further demonstrate the ability to identify single somas in sparsely labeled mouse cortex during both structural imaging of cells labeled with enhanced green fluorescent protein (eGFP) and functional imaging of GCaMP6s-expressing somatostatin (SST) cells.

## RESULTS

### CMOS Implantable Fluorescence Imager for Deep Neuronal Imaging

Acus has a shank form factor with two 4.1 mm-long 120 μm-wide shanks with a 180-μm separation between them (Fig. 1a). Each shank contains two rows of 128 single-photon avalanche photodiodes (SPADs) arranged with a longitudinal pitch of 25.67 μm (x-direction) and lateral pitch of 73.22 μm (y-direction). The 7.7-μm-diameter pixels yield a 6.3% fill factor, with the remainder of the area occupied by CMOS circuitry. Each pixel includes a quenching circuit to reset the SPAD, a six-bit counter to enable photon counting up to 63 photons per pixel, and pixel addressing logic. The base of the device contains a digital-to-time converter for on-chip clock generation and metal-oxide-metal capacitors to decouple the high voltages necessary for Geiger-mode operation of the SPAD pixels. Overall, the imager is able to record within a volume of approximately 3.4 mm × 600 μm × 200 μm in the longitudinal (x), lateral (y), and vertical (z) directions, respectively. The actual imaging volume is currently limited by the extent of excitation light delivery.

**Fig. 1.**
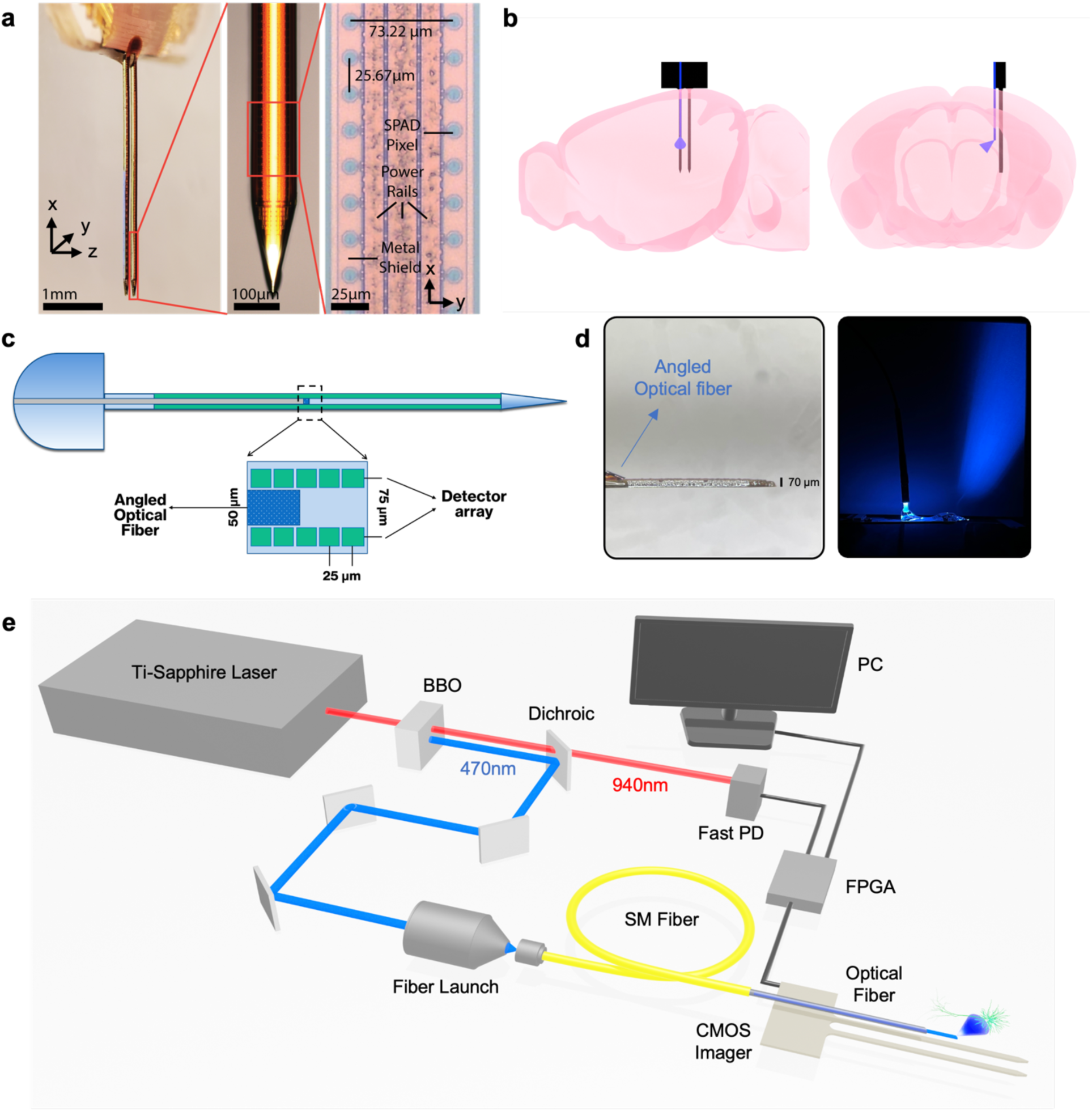
Acus system overview. **a**, The Acus CMOS implantable imager consisting of two 4.1-mm long, 120-μm wide shanks, each containing 2×256 SPAD pixels. **b**, Insertion of Acus into the mouse brain for deep neural imaging. Sagittal cut on the left, coronal cut on the right. **c**, Placement of the 50-µm optical fiber between the rows of SPAD detector array. Zoom-in version illustrates the tip of the fiber and the two-row pixel array. **d,** On the left, side view of the shank along with the co-integrated fiber is shown. The image on the right demonstrates the illumination of the fiber when activated in the dark. **e**, Optical setup where the excitation light source (470 nm) is generated from Ti-Sapphire laser (940 nm) and fed to the Acus through a single-mode fiber (SM Fiber). A fast photodiode (Fast PD) is used to generate clock signal from detected laser pulses for image acquisition.

In lieu of optoelectronic light sources on the shanks of Acus itself, our first implementation uses a 50-μm single-mode optical fiber which is co-packaged with the imaging shanks. The fiber has an aluminum mirror at its tip to provide right-angled illumination (Fig. 1b). The fiber light source is packaged (see Methods) such that it lies in the space between the two rows of SPAD pixels on each shank (Fig. 1c) with the light delivered at a 25° angle relative to the sensor plane at the distal end of the 70-μm shank (Fig. 1d). After the excitation light is launched from the fiber at this angle (Supplementary Fig. S1), due to scattering anisotropy (g=0.887), the direction of the excitation photons away from the shank is largely maintained, allowing this to deliver two orders of magnitude of additional background rejection, complementing that provided by the optical filters. In our *in vivo* studies, only one shank is packaged with the illumination fiber to reduce packaging complexity. The external light source coupled to this fiber is a titanium-doped sapphire (Ti-sapphire) solid state laser with a beta barium borate (BBO) crystal (Fig. 1e) to provide conversion of 940-nm infrared light to 470-nm blue light. On a separate optical path created by a dichroic mirror, a fast photodiode (PD) is used to sense laser pulses to generate a synchronizing clock signal. Total optical output power delivered with the fiber was measured to be approximately 200 µW, resulting in an optical output power density of 0.1 µW/µm^2^ from the fiber tip.

### Spectral-Temporal Filtering for On-Chip Fluorescence Imaging

Effective filtering of the excitation light is one of the most critical components in fluorescence imaging systems. In the absence of adequate filtering, the imaging sensor receives significant numbers of excitation photons, wasting available dynamic range (DR) of the imaging electronics. More importantly, however, these excitation photons add shot noise and degrade signal-to-noise-ratio (SNR) performance, limiting the sensitivity for detection.

The Acus device is fabricated in a commercial 130-nm high-voltage CMOS technology; custom post-processing is used to monolithically integrate spectral filters to provide rejection of the excitation light source as well as to shape the chips into a needle-like form factor (Fig. 2a and Methods). Final cross-section of the Acus shanks after the post-processing is shown in Fig. 2b. The first post-process fabrication step is the sputter deposition of a thin-film interference filters consisting of alternating layers of tantalum pentoxide (Ta_2_O_5_) and niobium pentoxide (Nb_2_O_5_). This filter provides greater than optical density (OD) 3 rejection at 470 nm while providing greater than 85% transmission at 520 nm for angles of incidence (AOI) between 0 and 45°. The effectiveness of this filter is reduced for photons coming from oblique angles as the spectral response shifts towards shorter wavelengths (Fig. 2c), requiring the addition of a dye-based absorption filter, as described below.

**Fig. 2.**
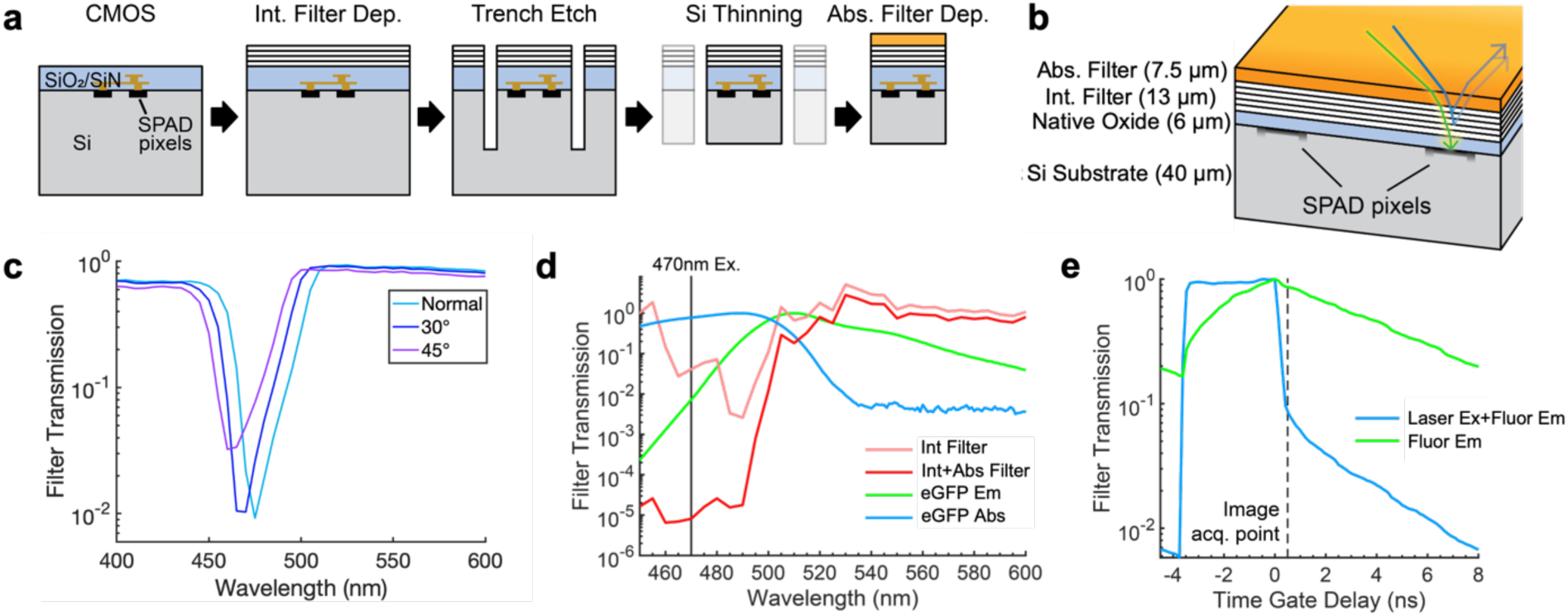
Post-processing of Acus into 70-μm-thick shank format with integrated emission filters. **a**, CMOS post processing sequence consisting of interference filter deposition, excimer-laser trench etching, silicon substrate thinning, and absorption filter deposition. **b**, Stack-up of post processed implantable imager with corresponding thickness of each layer. **c**, Interference filter transmission at different angles of incidence; 0° (normal), 30° and 45°. **d**, Transmission of the on-chip interference alone as a function of wavelength (“Int Filter”) and transmission of absorption-interference spectral filter (“Int+Abs Filter”). Also shown are the emission and absorption spectrum of eGFP (“eGFP Em” and “eGFP Abs”). **e**, Effective transmission of time-gated SPAD temporal filter; image acquisition starts after the laser pulse reaches 0.1 of its peak value. Starting the integration of fluorescence emission (“Fluor Em”) from this point adds an effective background rejection of OD 1 thanks to the fast SPAD impulse response.

After the interference filter deposition, a micromachining excimer laser is used to etch 100-μm deep trenches around the periphery of the active imaging area. Then, back-side milling reduces the die thickness to approximately 70 µm to release the shank from the rest of the die. Reducing the thickness further leads to warpage of the imager-filter complex due to the coefficient of temperature expansion (CTE) mismatch between the interference filter and the silicon substrate. As the last step of the post fabrication, a dye-based absorption filter (see Methods) is applied to the surface of the imager at a thickness of 7.5 µm which provides an additional OD 2.6 of rejection at 470-nm with respect to 80% transmission at 520 nm independent of AOI. The interference and absorption filter combination provides at least OD 5.6 rejection at 470 nm (Fig. 2d).

In addition to these spectral filters, the use of time-gated SPAD pixels allow for additional temporal filtering of the excitation light; that is, the SPAD pixels are gated based on the timing of the laser pulse, relying on the fluorescence lifetime to allow light to be collected after the excitation light has been removed (Fig. 2e). This gating provides an additional order of magnitude of background rejection, limited by the impulse response of the SPAD detectors, as shown in Supplementary Fig. S2c. The hybrid absorption-interference emission filter along with temporal time-gating provides a total background rejection of approximately OD 7.

Given that there is sufficient dynamic range to collect unrejected background light without saturation, the static background can be subtracted. However, the challenge is that the photon shot noise associated with this background cannot be eliminated, and it degrades the achievable signal-to-noise-ratio (SNR) for fluorescence detection. Fig. 3a illustrates how the observed rejection improves with the addition of the interference filter, absorption filter, and temporal filter at an excitation power of 0.2 nW/µm^2^ delivered uniformly through an objective from the top. To better quantify the effects of unrejected background on SNR, we use Acus to capture images of a single fluorescent 10-μm-diameter microsphere (see Methods) with a fluorescence emission spectrum similar to eGFP at approximately four times the brightness of a labeled neuron. This microsphere is positioned 150 μm above the Acus shank. As we increase the background rejection with each subsequent filter addition, we observe an increase in the observed SNR, achieving approximately 33 dB with full filtering.

**Fig. 3.**
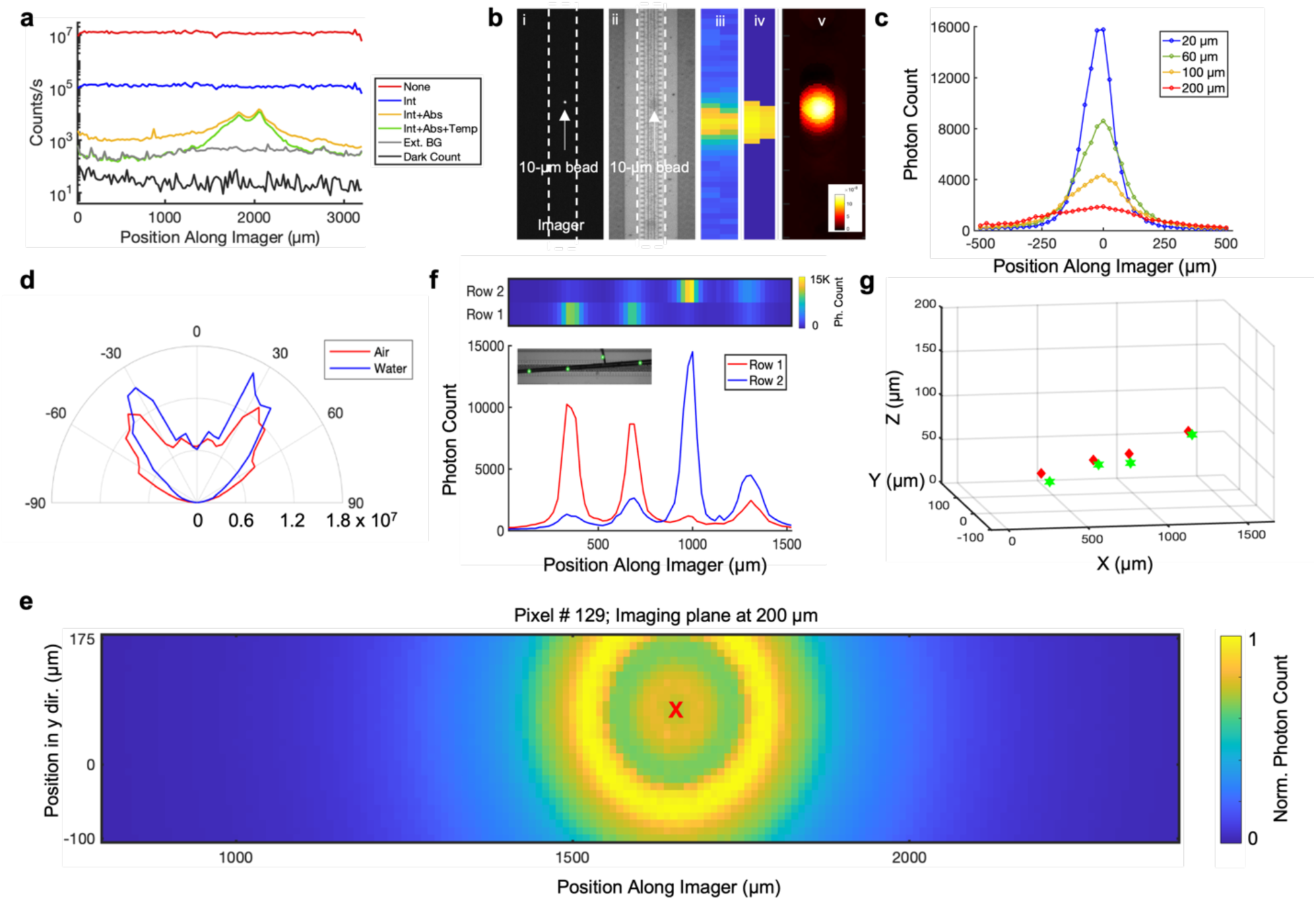
SPAD imager characterization. **a,** Contributions of different levels of spectral and temporal filtering of background excitation light while imaging a single 10-µm fluorescent microsphere: no filtering (“None”), interference filter alone (“Int”), combined interference-absorption filter (“Int+Abs”), combined interference-absorption filter with temporal time-gating filter (“Int+Abs+Temp”). “Ext. BG” denotes the excitation background in the presence of full filtering while “Dark Count” denotes the DCR. **b,** Raw image of a single 10-µm microsphere at a z-distance of 150 µm above the imager: (i) confocal fluorescent image, (ii) confocal widefield image, (iii) raw image from Acus, (iv) simulated raw image, (v) reconstructed point-source from BSS analysis of the raw image. **c,** Measurements of single 10-µm fluorescent microsphere at different z-distances, showing the degradation of resolution as the distance increases. **d,** Angular profile of the Acus SPAD pixels in two different media: air and water. **e,** Collection field of a single SPAD pixel (#129) and its normalized transmission function at an imaging plane 200 μm above the imager array. Pixel location is indicated with the red “X”. **f,** Image of four 10-µm fluorescent microspheres as a function of position along the imager. The color bar of the top plot shows the range of photon counts from 0 to 15000. Bottom plot shows response of each row on Acus separately. Inset shows the confocal image of the microspheres attached to custom-made pipettes. **g,** Volumetric localization of these four microspheres above Acus through BSS. Red and green markers show the true locations and estimated locations of the microspheres, respectively.

### Blind Source Separation with Angle-Sensitive SPAD Pixels

In Fig. 3b, with the same microsphere as in Fig. 3a, we present a reconstructed planar image acquired by the pixel array with full filtering, which corresponds closely with that taken by a widefield confocal microscope. In Fig. 3c, we position the microsphere just 20 μm above the array and then gradually move it away from the imaging plane, resulting in decreasing resolution, as measured by an increase in the full-width-at-half-maximum (FWHM) of the sphere in the raw image as the distance to Acus increases (Supplementary Fig. S3 & S4), exactly as expected with a lensless contact imager in the absence of an amplitude or phase mask.

Lensless imagers introduce such masks between the scene and the imaging device^28–32^ to perform inverse imaging^33^, that is, to obtain an image as a function of spatial coordinates. While this is well suited for mesoscopic imaging of population dynamics for fluorescently labeled neurons, if single-unit fluorescent recording is desired, source separation and localization is a far more effective approach to extracting information from the scene. These approaches are further enabled by an emerging class of fluorescent reporters that are soma-localized^34–36^, providing “point-source-like” structure to the spatial distribution of fluorescence in a neuronal population.

Blind source separation (BSS) methods can extract these fluorescent signals from mixtures of many of these sources fluorescing simultaneously.^37^ BSS is generally a difficult problem to solve; it relies on the innate properties of the signals making the mixture: sparsity, synchrony, temporal structure and amplitude distribution. In our case, in addition to source separation, we also need to perform source localization, the problem of assigning a spatial location to a signal extracted by source separation. To enable both BSS and localization, Acus must provide both a sufficient number of detectors (which must significantly exceed the number of neurons to be identified) and spatial selectivity, meaning that the collection fields for each of the pixels on the device have sufficient spatial diversity to allow BSS methods to perform the separation. Here, collection field refers to the characteristics of the solid-angles of collection for each pixel.

The collection fields for the pixels of Acus are defined by the filter stack and back-end CMOS, resulting in the angular profile for the light transmission shown in Fig. 3d at 520-nm wavelength.^28^ This collection field is manifested in the halo-like features in the spatial sensitivity of an individual pixel as demonstrated in Fig. 3e at a distance of 200 μm. The refractive index of the medium plays a crucial role in this angular dependency; in particular, the angular profile gets narrower as the refractive index of the medium increases (Fig. 3d).

We developed a customized BSS algorithm with multi-step sparsity-regularization based on the characteristics of this pixel collection field, allowing us to localize sources in three-dimensional (3D) space (see Methods). With this algorithm, high-dynamic-range signals with high-spatial-frequency content, characteristic of Acus, are used to reconstruct a sparse scene of neuronal point sources. Fig. 3f shows the raw image of four 10-μm fluorescent microspheres hovering above the imager. The result of the source separation algorithm in localizing these sources in 3D volume is presented in Fig. 3g. This volumetric imaging can be performed on Acus at frame rates as high as 400 frames/sec, which will ultimately be necessary for the detection of recently developed GEVIs.^38–40^ Supplementary Movies 1-6 show the experiments where the localization of a single 10-μm fluorescent source moving in 3D space at various frame rates is performed.

### Single-Neuron Imaging with Acus

Calculations indicate that the effective OD-7 filtering on Acus should be sufficient to resolve a single eGFP-labeled neuron at distances of up to 140 μm (Supplementary Fig. S5) at a light intensity of 0.1 μW/μm^2^ from a 1 μm^2^ source which is assumed to be emitted at the plane of the detector with a Gaussian profile with 8° half-angle. This can go up to 170 µm if the excitation power is increased to 1 μW/μm^2^. In general, the SNR increases as the square-root of the excitation power with this power level, although ultimately being limited by tissue heating.

To verify this experimentally, we first perform imaging on 100-µm slices extracted from the cortex of an eGFP-labeled mouse with sparse neuronal expression (see Methods). In these experiments and the following *in vivo* experiments, excitation is delivered with a 50-µm optical fiber, positioned in an angled manner for the maximum power delivery to the neuron while minimizing the collection of background excitation light. Monte Carlo simulations of the resulting excitation power (Supplementary Fig. S1) indicate that an optical power of 200 µW at the fiber output leads to 12 µW power delivery to a 15-µm-diameter neuron.

Even though expression is not strictly limited to the soma, we observe that the somas have brightness significantly above that of the neuropil in this eGFP labeling. We place the slices on top of Acus, which allows a confocal microscope to provide the ground truth location for the single neuron within the slice (Fig. 4a). One such neuron is located at a position 130 μm above Acus as shown in the raw image traces of Fig. 4b. BSS analysis of this detector data localizes the neuron at the position shown in Fig. 4c, which matches the position determined by the confocal microscope.

**Fig. 4.**
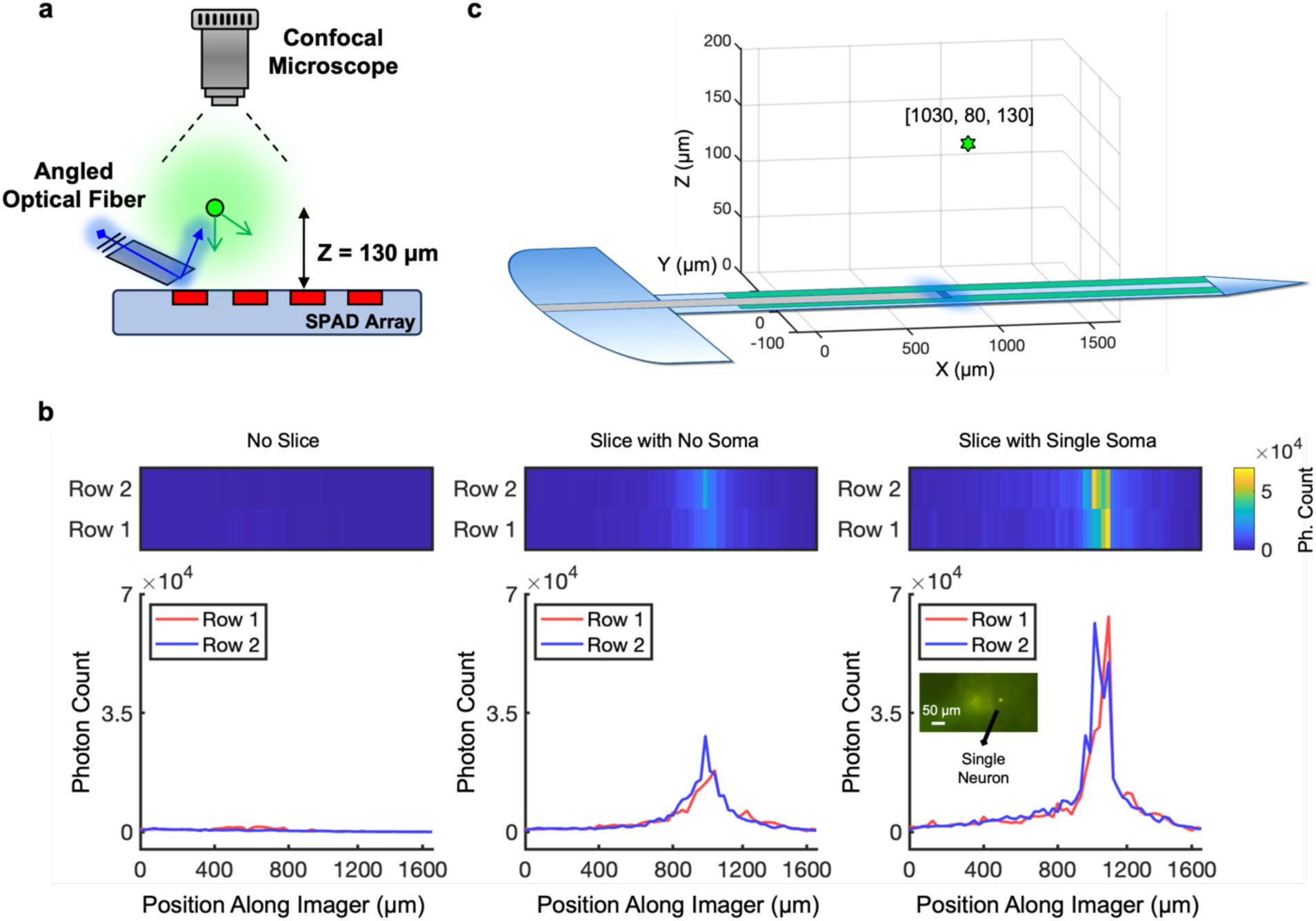
Structural imaging using Acus in 100-µm mouse brain slices. **a,** The arrangement of the *in vitro* brain slice experiments. The slices are concurrently imaged by Acus at the bottom and by a confocal microscope at the top. **b,** Images acquired by Acus for an integration time of 62.5 ms as a function of position along the imager for three cases: no slice present (“No Slice”), slice with only neuropils and no soma present (“Slice with No Soma”) and a slice with detected neural soma (“Slice with Single Soma”). Photon count observed for the “Slice with No Soma” reflects the background over the localized excitation region of the optical fiber, consisting of unrejected excitation and emission resulting from neuropils. Inset of the “Slice with Single Soma” plot shows the confocal image of the neuron as ground truth. **c,** Estimated location of the imaged neuron in 3D volume with BSS.

### *In vivo* BSS

Following the characterization of a single neuron in a brain slice, we proceed to assess Acus’ ability to detect single neurons in *in vivo* setting in the same eGFP-expressing mice. To achieve this, we perform a cranial window and insert Acus into the cortex at a rate of 50 μm/sec, collecting data at a frame rate of 40 frames/sec during the insertion process as illustrated in Fig. 5a. Three raw images captured at different time points during the insertion are displayed in Fig. 5b (Supplementary Movies 7 and 8). A single labelled neuron is captured by Acus, and BSS analysis of the acquired images accurately records the relative position of the neuron as the shank approaches to the neuron during insertion. (Fig. 5c). These experiments were repeated on two different eGFP-expressing mice (n=2).

**Fig. 5.**
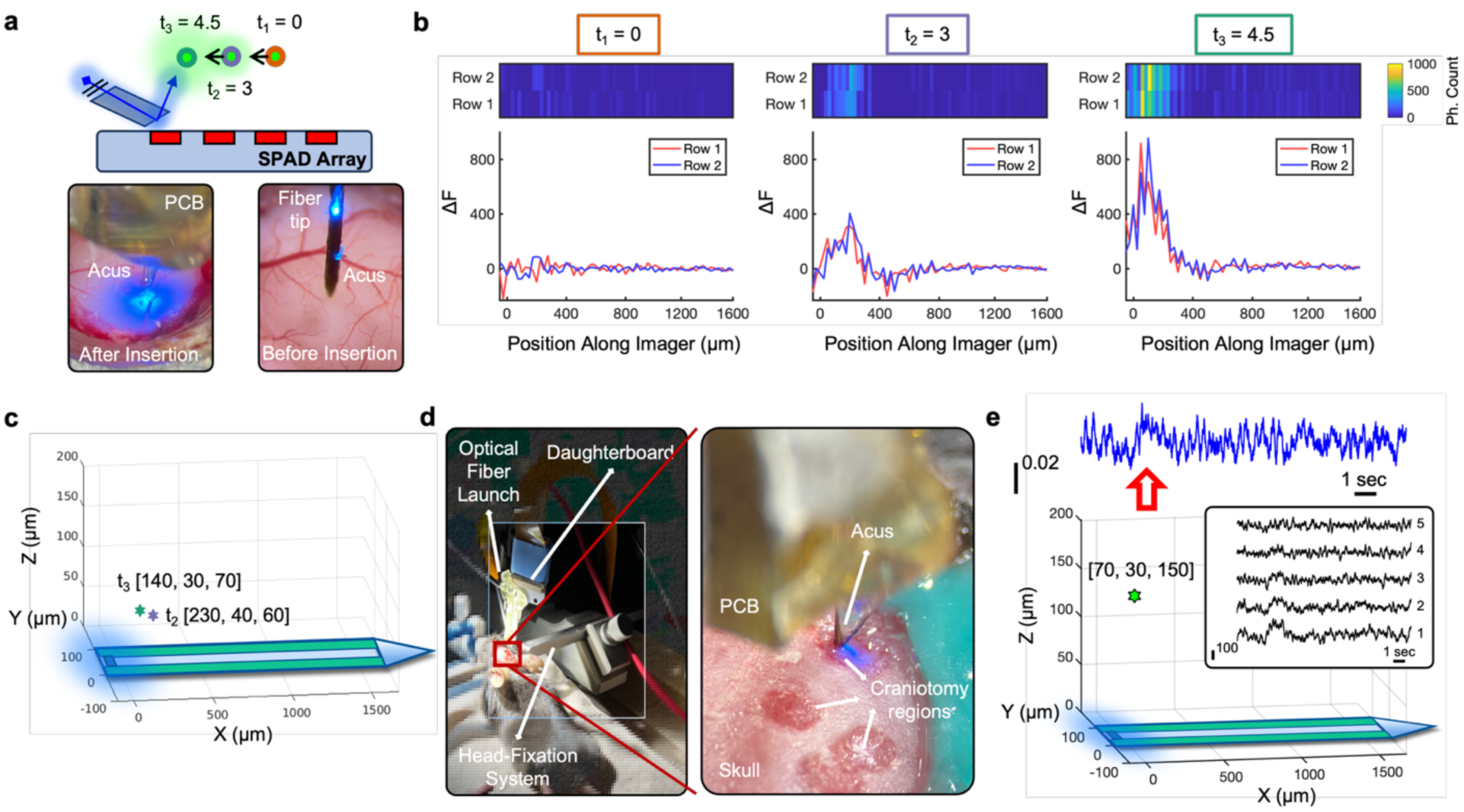
*In-vivo* lens-less imaging. **a,** Single neuronal imaging during insertion of Acus into brain tissue at a rate of 50 μm/sec. Three distinct time points during insertion are illustrated, namely t_1_, t_2_ and t_3_. Inset images show the insertion procedure with two images taken before and after insertion of Acus into the mouse cortex. **b,** Images acquired by Acus at an integration time of 25 ms as a function of position along the imager for the three time points shown in (a): t_1_=0 (orange) -the starting point-, t_2_=3 seconds (purple) and t_3_=4.5 seconds (green) after the starting point. **c**, Estimated location of the neuron using BSS analysis at t_2_ and t_3_. **d,** Implantation of Acus into the mice cortex through a small craniotomy opening for functional GCaMP6s imaging. **e,** Estimated location of the detected neuron in 3D volume as determined by BSS analysis. The inset shows photon count acquired by Acus at 400 frames/sec moving along the shank in the direction of increasing *x*. Each trace is the summation of 16 pixels (block of 2×8). In the trace labels, “1” denotes the block extending from *x* = 0 to 200 μm, with “5” denoting the block extending from *x* = 800 to 1000 μm. Data are detrended and processed with a moving average filter prior to summation. The top plot shows the backpropagated signal at the voxel site [70, 30, 150]. Red arrow indicates the onset of the calcium activity.

Acus is also able to detect functional single-unit activity in animals with sparse GCaMP6s expression.^41,42^ These animals are bred through Cre-lox system (see Methods), resulting in the selective expression of GCaMP6s in somatostatin-expressing neurons (SST cells). Fig. 5d shows the experimental setup for the *in vivo* study after we inserted the Acus into the mouse cortex and positioned it near an SST cell. Measured time-domain data showing the spontaneous calcium activity of a single SST cell as measured by detectors along the Acus shank and the estimated location of the cell resulting from the BSS analysis are both shown in Fig. 5e (Supplementary Movie 9). Channel groups are independently rescaled to have the same noise level, to account for non-uniform excitation light (Supplementary Fig. S1). Fig. 5e also shows the time-domain trace backpropagated to the localized voxel at which the neuron is detected (see Methods).

Extended Data Fig. 1a-b shows results from another single neuron recording in a second experiment in a different animal. In this case, we placed a tungsten electrode 50 μm from the surface of Acus and apply electrical stimulation pulses (100 µA, biphasic, 100-μs pulse widths for both anodic and cathodic phases, 10 pulses, 500-Hz repetition rate). Measured time-domain fluorescence signals are shown in the presence and absence of electrical stimulation as measured by detectors along the Acus shank. In Extended Data Fig. 1c we show an average waveform derived from multiple (n=97) stimulated single-neuron responses measured from 33 different neurons, which shows the characteristic GCaMP6s waveform shape.^43,44^

To demonstrate the high-frame-rate imaging capabilities of the device, we inserted Acus into the cortex of a densely labeled GCaMP6f mouse.^42,43,45^ In this experiment, a tungsten electrode was also positioned to perform electrical stimulation with the same protocol used for the single-neuron experiments of Extended Data Fig. 1. Acus measures the calcium signal evoked by the electrical stimulation of a large population of neurons, which falls off with distance from the stimulation site (Extended Data Fig. 2). Imaging at 400 frames/sec allows precise capture of the approximately 35-ms rise-time-to-peak of the GCaMP6f signal. In an experiment performed in a different animal (n=2), Supplementary Fig. S6 provides an example of detected spontaneous activity where there was no electrical micro-stimulation applied. Subsequently, control experiments with wild-type animals have been performed to confirm that the source of the signal acquired by Acus is only from fluorescent GCaMP activity (Supplementary Fig. S7a) and that the switching of the excitation light does not cause any neural response (Supplementary Fig. S7b).

To study tissue heating during illumination, we analyze the heating distribution through the use of a FLIR infrared camera (Supplementary Section S1 and Supplementary Fig. S8c). Experiments performed with 100-µm brain slice show that temperature variations are less than 1°C for optical output powers of 200 µW at the output of the fiber (Supplementary Fig. S8b). We were able to perform *in vivo* experiments that lasted more than four hours with Acus without causing any damage to the tissue or showing any decline in the performance or image quality, demonstrating its suitability for studies requiring measurements taken over extended periods.

## CONCLUSIONS

In this work, we have presented Acus, an implantable CMOS optical recording system for fluorescent interrogation of live neural tissue at depth. The key innovations are a shank-based implantable imager design incorporating SPAD detectors and integrated electronics, that mirrors CMOS electrophysiological probes. For soma-localized fluorescent reporters, BSS algorithms are used to determine the location of individual labelled neurons in three dimensions. Integrated spectral filters on the shank, augmented by time-gated temporal filtering, reject background excitation light, providing enough SNR performance for both structural and functional imaging of a single neuron in live mouse brain. The use of SPAD pixels provides advantages over traditional photodiodes, including enhanced sensitivity and dynamic range and narrower impulse response functions.^46–48^ Our device has the ability to image at frame rates up to 400 frames/sec, a necessary requirement for emerging soma-targeted voltage reporters with cell-type specific expression.

There are several notable limitations with the current implementation of Acus. Acus currently uses a fiber for light delivery, which significantly limits the field-of-view for imaging and results in increased tissue damage on insertion. In future versions, micro light-emitting diodes (μLEDs) will be integrated directly onto the shank. With recent laser ablation methods developed for ultra-thin μLED production and further back-side thinning of the silicon, the total thickness of the μLED-integrated imager could be made as low as 40 μm, which paves the way for multiple-shank systems to image larger brain volumes.^49–51^ Furthermore, novel SPAD designs will enable tighter pixel pitch and improved fill-factor, supporting larger numbers of detectors in the same area with improved collection efficiency. Phase-modulating optical metasurface designs optimized for the BSS algorithms will allow for better demixing of individual labeled neurons while maximizing the photon collection.^52–54^

## METHODS

### Shank Form Factor and Silicon Post-Processing

The CMOS substrate for Acus is designed in a 130-nm CMOS process (Taiwan Semiconductor Manufacturing Company, Hsinchu City, Taiwan). After commercial fabrication, additional post-processing steps are performed. First, we define the shape of the insertable shank by trenching the surrounding silicon by means of two methods. First method is thermal ablation with a Cu-vapor micromachining laser (IPG Photonics, Oxford, MA, USA) which has a wavelength of 532 nm.^55^ 100-µm-deep trenches can be created when a power of the device is set to 80% with a repetition rate of 120 kHz, pulse length of 10 µs and 201 passes are repeated. Second method to define the trenches along the imager is deep reactive ion etching as described in Supplementary Section S2.

After trenching, the die is thinned with an XPREP mechanical polisher (Allied High Tech, Compton, CA, USA) to a thickness of 40-70 µm, releasing the probes from the die (Supplementary Fig. S9 & S10). Supplementary Fig. S11 shows that there is no degradation in performance after the post-processing. The chips are subsequently wirebonded to the customized daughterboard printed circuit board (PCB) with a 51-pin connector. This daughterboard PCB connects to a motherboard to allow data transfer to support a maximum frame rate of 51,000 frames/second.^29^

An externally provided global *LASER_IN* shutter clock allows time-gating of the detectors to be synchronized with excitation laser pulses at 80 MHz. Custom MATLAB code is used to program the phase and duty cycle of this gating to prevent the on-chip counters from saturating.

### External Electronics

The daughterboard is connected through a flexible cable to a motherboard, carrying an Opal Kelly XEM6310, voltage regulators and decoupling capacitors for each voltage line. A Xilinx Spartan 6 FPGA on the Opal Kelly is used to implement a state machine that controls communication and clocking of the system. Duty cycle, phase, and the number of frames to be captured for a given configuration are all programmed into the state machine. This controller cycles through the 512 pixel addresses to program an enable signal into each pixel’s local one-bit memory, a process that takes approximately 25 µs. The global SPAD ON clock activates all SPADs that were thus enabled.

### SPAD Operation

Supplementary Fig. S12a shows the operation of a SPAD pixel with time-gating and in-pixel quench-and-reset circuitry.^29^ A given SPAD is reverse-biased beyond the breakdown voltage to enable it for detection. This reverse bias for each of the 512 SPADs is established between a static global cathode voltage at 17 V and a dynamic anode voltage (*V_ANODE_*) which is independent for each pixel and exceeds the breakdown voltage (*V_BD_*) of the SPADs measured as 15.5 V. When *Vanode* is biased at ground, the reverse bias voltage across the SPAD becomes equal to the cathode bias, putting the SPAD into Geiger mode and enabling it for photon detection. When *Vanode* is at the positive reset voltage (*Vrst*), the voltage across the device is lower than *Vbd*, and the SPAD is disabled and insensitive to incoming photons. The rising edge of the *ON* signal globally triggers this switching of *Vanode*.

In pixel operation, the large reset NMOS transistor (M1) begins discharging anode until *V_ANODE_* drops below the threshold (∼750 mV) of the inverter acting as a comparator. When the inverter output goes high, the flip-flop edge detector is reset, turning M1 off and setting anode to a high-impedance state. The SPAD is now biased in Geiger mode and is ready to detect a photon. The long-channel half-latch transistor (M2) provides resistive path at the anode terminal of the SPAD. During an avalanche breakdown event, the resulting current flow causes the voltage at the input of the comparator to increase, flipping the comparator, triggering the event detection flip-flop, and incrementing the counter by one. The flip-flop ensures that events are only detected when *ON* is enabled. To suppress afterpulsing, the counter is gated such that at most one photon can be detected per laser cycle. When the counter reaches 63 counts, the *Full* signal is asserted and the counter stops incrementing.

The timing diagram in Supplementary Fig. S12b shows typical SPAD operation in a cycle in which a photon is detected and in a cycle in which no photon is detected. In typical *in vivo* experiments, fluorescence expression levels and excitation intensities generally result in a probability of detecting a single photon in a given laser pulse of only 7.25%. Each laser pulse induces an exponentially decaying fluorescence emission determined by the lifetime of the fluorophore. During the first pulse in Supplementary Fig. S12b, no photons arrive at the device, and *V_ANODE_* returns to *V_RST_* without incrementing the *Count* signal. During the next laser pulse, a single photon arrives at the device, producing an earlier rise in *V_ANODE_* and “quenching” the avalanche current through self-timed feedback. The event detection circuit block increments *Count* by one.

### SPAD Characterization

Key performance metrics of the P-type (N-implant) shallow-junction SPAD, manufactured in the 130-nm CMOS process, were characterized in an integrating sphere and published in a previous publication.^29^ On average, approximately 2% of the SPADs are measured to be “hot” with a dark count rate (DCR) that is five standard deviations away from the mean DCR of 40 Hz for a *V_ov_* of 1.0 V. At this overdrive voltage, the median photon detection probability (PDP) of the SPADs in the Acus device is 10% for 520-nm fluorescence light emitted by eGFP.

A useful figure-of-merit (FoM) for chosing the optimal value of *Vov* in the case of fluorescence imaging is given by:

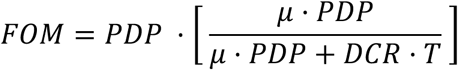

where *μ* is the mean value of the fluorescence photon emission rate dictated by a non-homogenous Poisson process and T is the measurement window length.^56^ Acus SPADs are able to achieve a score of 0.92 on average with this FoM a fluence of 10^5^ photons per second, equivalent to a photon flux of 40 fW/m^2^ at a 520-nm wavelength. Due to the low DCR relative to fluence, the optimal operating value of *V_OV_* across this range is found to be 1.0 V. Operation at this relative low overdrive voltage also has the benefit of also reducing afterpulsing events.^57^ We can characterize this afterpulsing by measuring the changes in DCR as the active-reset dead time is swept from 300 ns to 10 ns. The nearly constant DCR over this range confirms that afterpulsing is not a significant contributor to DCR.^58^ Acus SPADs show high linearity for a wide dynamic range where the linearity is limited by the dark count at low fluence and by photon pile-up at high fluence.

Supplementary Fig. S8a shows the DCR measured for different temperatures. The median DCR increases by approximately 60% when the temperature is increased from 30°C to 40°C, comparable to the difference in temperature between typical room temperature and the ambient temperature in the mouse brain.^58^

Supplementary Section S3 and S4 presents more information on the characterization of the imager array and Supplementary Table S1 compares Acus with other time-gated SPAD imagers. While much of the novelty of this work lies in the form factor and its packaging with spectral filters, pixel performance compares favorably with other time-gated SPAD imagers in conventional planar array formats where area and power are not as limited.

### Time-Gate Filter Operation

Since there is no substantial difference in PDP between excitation and emission wavelengths, spectral filters must be incorporated to block excitation light. For a standard free-space epifluorescence microscope, a “filter cube,” consisting of an emission filter, an excitation filter and a dichroic mirror, provides OD > 6 rejection of the excitation wavelength. Achieving the same levels of rejection with an integrated filter on Acus remains challenging; thin-film interference filters, providing the best filter contrast, do so for light that is not impinging at oblique angles (Supplementary Section S5). To extend reduction to this oblique-incidence light, these filters must be supplemented with effective dye-based absorption filters. Absorption filter applied on Acus consists of 95 mg Valifast Yellow 3150 (Orient Corporation of America) mixed with 200 µL KMPR 1000 (Kayaku Advanced Materials, Inc.) and 600 µL cyclopentanone (Supplementary Fig. S2 & S13).

Time-gating provide an additional degree of rejection beyond spectral filtering. Fluorophores display an exponentially decaying fluorescence intensity after pulsed excitation with a decay time given by the fluorescent lifetime. Typical lifetimes are on the order of nanoseconds (e. g., 4.1 ns for eGFP in phosphate buffer solution, 4.0 ns for fluorescein, and 1.68 ns for Rhodamine B). Using a pulsed excitation source and having the ON signal go high a short delay (< 1 ns) after the end of the laser pulse, SPAD detection can be gated on after the excitation pulse has ended but fluorescence persists. Time-gating performance for a single 10-µm fluorescent microsphere is presented in Supplementary Fig. S2d.

### Blind Source Separation Algorithm for High-Frequency Signals with High Dynamic Range

The detectors in Acus have an angular sensitivity determined by both the filter stack and the back- end metal of the CMOS process (Supplementary Fig. S3 & S14). This angular sensitivity combined with the SPAD configuration creates a detection field above Acus covering a volume of 3.4 mm × 600 μm × 200 μm, consisting of 408,000 cubic voxels with dimensions 10 μm × 10 μm × 10 μm. With 512 total detectors in Acus, the detection field (***A***) has dimensions 512×408,000, in which each column is a normalized raw image of a single 10-μm fluorescent source in a given voxel.

The measured 512×1 raw image, **y**(t), is given by ***y***(t) = ***A*** ∗ **x**(t) + ɛ, where ɛ is the 512×1 shot noise vector for a ΔF image (F-F_0_, where F_0_ is the average of all frames in that measurement session) and **x**(t) is a 408,000×1 binary vector representation of all the voxels of the imaged volume sampled compressively through the detection field ***A***. Each element ***A****_ij_* represents the response of pixel *i* to voxel *j*. Each row of ***A*** captures the angularly modulated detection field of each pixel as shown in Fig. 3d.

Pseudoinverse backprojection of a single-voxel source by the least-mean-squares error solution is used to calculate the point spread function (PSF) of the system:

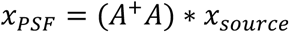

where ***A^+^*** is the left pseudoinverse of ***A*** and *x*_*source*_ is a 408,000×1 vector in which all elements are zero except one. A sample simulated PSF is displayed in Supplementary Fig. S15a-d for a point source located at [550, 90, 50]. The FWHM of the PSF for each voxel can be calculated from Gaussian fits. Supplementary Fig. S15e shows the resulting FWHMs for voxels of different z heights, showing the diminishing resolution as the fluorescence source moves away from the imager plane.

From sparse reconstruction theory, we know that to localize a given number of sources, we need at least as many recording sites. We perform maximum likelihood estimation to determine the location of the (small number of) signal sources even though the number of voxels, and thus the dimensionality of the problem, is much higher in this underdetermined system.

With multiple fluorescent sources in a scattering and absorbing volume, Acus obtains signals with very high dynamic range; signals from more distant neurons are substantially weaker than those from neurons close to the device (Supplementary Fig. S16). In addition, the skewed aspect ratio of the detector array combined with the angle-sensitivity of the pixels causes sources close to the array to produce significant high-spatial-frequency components in the raw image, manifesting as low-magnitude singular values of the mixing matrix ***A***. Moreover, tissue scattering and the spectral spread of emission impairs the effective resolution of angle-sensitive pixel array. These high-spatial-frequency, high-dynamic-range raw signals are poorly served by conventional penalized regression models for sparse selection, such as Lasso and Ridge regression, which are overwhelmed by neurons close to the imager and are completely unable to detect those further away. Instead, we develop a BSS algorithm based on a least-squares solver with multi-step sparsity-regularization which performs well on the type of data collected by Acus. This algorithm decouples the signal separation and regression analysis to extract valuable information from the y-axis measurements while maintaining the same high-quality signal separation as traditional line imaging methods. This strategic approach allows us to effectively use spatial information across all three dimensions, resulting in improved signal resolution and fidelity.

Raw images are captured from Acus as photon counts as a function of the corresponding pixel address (Supplementary Fig. S17). The first step of data processing is to remove hot pixels from the images. Hot pixels, which occur at an incidence of approximately 2% in our process, are characterized by anomalously high DCR and result from the presence of traps in the vicinity of the multiplication region of the SPAD. Pixels with DCR more than five times the mean are identified as hot, and their photon count is replaced by the mean of the adjacent pixels.

In order to determine the number of sources present in the scene, we need to establish a mask to identify regions in the raw image corresponding to the response of different neurons. This mask allows the individual neural responses to be normalized and localized one-by-one. To establish this mask, the raw images are processed with a moving average filter to remove high-spatial-frequency components beyond 50 µm^−1^, which is accomplished with a 2×2-pixel moving average. The resulting averaged image, normalized to the highest photon count, shows distinct peaks. (Supplementary Fig. S17b). Each peak in the averaged image is identified by its prominence, which is defined as the minimum vertical distance that the signal must descend from the peak on either side before either increasing again to a level higher than the peak or reaching an endpoint. The threshold prominence in our analysis is determined to be 0.1. Peaks higher than a threshold prominence level of 0.1 are resolved as belonging to separate sources. The locations and widths of these peaks are then used to create masks which are applied to the original raw signal to extract the components of the original image associated with each independent source (Supplementary Fig. S17c). By this method, we are able to separate the signals resulting from multiple neurons in the 3D volume above Acus.

For each separated signal, regression is then performed by minimizing the L2-norm of the residual error with the constraint that the number of sources in the scene be one:

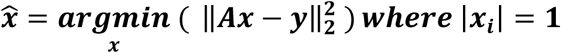

The residual error for the peak of highest prominence must be less than 1.5 in order for a fluorescent source to be identified. To avoid overfitting, *N* voxels with the highest maximum likelihood estimation are identified with *N* as a tunable parameter. The geometric center of the resulting N-voxel cluster is then used to localize the separated individual fluorescent source.

Presented BSS algorithm has been validated with 10-µm microspheres hovering above the imager in 3D space (Supplementary Fig. S18). The translational movement of individual microspheres has been facilitated with custom pipettes controlled with micromanipulators.

### Scanning Electron Microscopy Characterization

In order to verify the coverage and topology of the deposited optical filters, cross-sections of the shanks of the Acus devices were prepared for scanning electron microscopy (SEM) using focused ion beam (FIB) milling with a Ga ion source at an accelerating voltage of 30 kV and a probe current of 65 nA, followed by final polishing at 2.5 nA (FEI Helios Nanolab 660 DualBeam) (Supplementary Fig. S9, S10, and S13).

### Surgery and In Vivo Experiments

The Institutional Animal Care and Use Committee (IACUC) reviewed and approved protocols for Columbia University’s program for the humane care and use of animals and inspects the animal facilities and investigator laboratories. Evaluation of the implanted devices was performed in compliance with Animal Welfare and Columbia’s IACUC regulations under approved IACUC protocol AC-AABU1663 “Development of high-density, implantable recording, imaging and stimulating arrays”. For this project, three separate lines of mice were used for GCaMP imaging and eGFP imaging. For sparse GCaMP6s imaging, a mouse pair with strain Sst-IRES-Cre and strain Ai162(TIT2L-GC6s-ICL-tTA2)-D were acquired from Jackson Laboratory. Mice were bred to generate double heterozygous pups expressing GCaMP6s in SST interneurons. For population activity imaging with GCaMP6f, a mouse pair with strain Vglut1-IRES2-Cre-D and strain Ai148(TIT2L-GC6f-ICL-tTA2)-D were acquired from Jackson Laboratory. Mice were bred to generate double heterozygous pups expressing GCaMP6f in pyramidal neurons. Six-week-old eGFP mice with strain Tg(Thy1-EGFP)MJrs/J were also acquired from Jackson Laboratory that exhibit sparse neuronal expression for structural single cell imaging.

In preparation for the slice experiments, the eGFP-expressing mice were used. These mice were anesthetized using a mixture of ketamine and xylazine. Subsequently, transcardiac perfusion with 4% paraformaldehyde (PFA) was carried out. After perfusion, the mouse was decapitated, and the brain was carefully extracted. The extracted brain was then sliced into thin sections using a Compresstome VF 310-0Z. These sliced brain sections, measuring 100 µm in thickness, were immersed in 10% PFA for a duration of 24 hours. Following fixation, the 100-µm-thick brain slices were attached to horizontally positioned shanks on a carrier substrate as part of the experimental setup.

Acus achieves the proximate excitation light delivery to the neural tissue with a 50-µm optical fiber whose tip is cut with 45° angle and coated with an aluminum mirror to allow for orthogonal light delivery. Backside of the mirror is further coated with black paint to minimize the leakage of excitation light into the pixels (Supplementary Fig. S19). The fiber is placed on top of the CMOS imager array, between the two SPAD rows with an angle of 25° towards the distal end of Acus. (Fig. 1d) This optimum configuration maximizes the SNR by delivering high power to the neural tissue, while keeping the direct detection of excitation light low. The power density used during these slice experiments is kept at 0.1 µW/µm^2^ by considering the following *in vivo* experiments during which it is crucial to minimize tissue heating. The images were captured simultaneously using a confocal microscope with an objective positioned at the top (Fig. 4a). The slices were arranged in such a manner that sparsely distributed cell bodies (soma) were precisely positioned over Acus, while remaining within the observation field of the confocal microscope. This arrangement allowed for the simultaneous imaging of the same neuron by both setups, providing a reliable ground truth reference.

For *in vivo* experiments, mice were anesthetized using isoflurane, with an induction level set at 3%, while being provided with oxygen at a rate of two liters per minute. Physiological parameters were continually monitored and maintained at a consistent level throughout the experiments. First, scalp overlaying the implantation area was removed following the subcutaneous injection of 0.2 ml of bupivacaine. A custom-made head-plate featuring a circular opening at its center measuring 8 mm in diameter was securely attached to the skull using dental cement. Following this, a craniotomy measuring 3×3 mm was performed over the somatosensory cortex to expose the brain (Supplementary Fig. S6 and S20). The dura mater was then carefully removed using a fine-tip scalpel. The CMOS shank, co-packaged alongside an optical fiber giving 200 µW power (0.1 µW/µm^2^), was fixed to the arm of the stereotactic frame and positioned in a vertical orientation above the exposed region of the mouse brain. Subsequently, the shank was carefully inserted into the brain tissue. Due to brain wide expression in our animal models, we did not target a specific location but instead broadly implanted the probe within the somatosensory cortex. To ensure precise placement within the designated area, we used a micromanipulator with micrometer precision for implantation, following these coordinates relative to bregma: anterior-posterior: −1.2 mm to 1.5 mm, medial-lateral: 2.2 mm to 2.7 mm, with a depth 1 mm to 1.5 mm.

### Structural imaging

For structural imaging, eGFP mice were used and Acus is inserted into the brain at a constant rate of 50 µm/s. To precisely map the insertion site, continuous image capture was carried out at a frame rate of 40 Hz (Supplementary Movie 7) and 400 Hz (Supplementary Movie 8) throughout the insertion process. At the end of the recording session, the animals were sacrificed by cervical dislocation.

### Functional imaging

For functional imaging, sparse GCaMP6s expression was used for single-neuron imaging and dense GCaMP6f was used for the imaging of population dynamics with rapid kinetics. To induce specific neuronal excitation in the targeted area during *in vivo* functional imaging experiments, a commercial tungsten electrode (Microprobes for Life Sciences, Gaithersburg, MD) was positioned using a zero-drift micromanipulator (Sensapex, Oulu, Finland) 50-µm away from the Acus shank. Stimulation consisted of a train of 10 pulses of amplitude 100 μA^59^ with anodic and cathodic durations of 100 μs and a pulse repetition rate of 500 Hz. Stimulation pulses were generated using an Intan RHS2000 electrophysiology amplifier, synchronized with the imaging setup. All images were acquired at a frame rate of 400 frames/sec. Temporal channel data are detrended with a third-order polynomial fit over 15 seconds. For GCaMP6s, data are filtered with a moving average filter (window size of 50 frames) to remove high-frequency noise (-3-dB cutoff at 3.54 Hz). For GCaMP6f, data are filters with a fourth-order Butterworth filter with a −3-dB cutoff at 25 Hz. To account for the non-uniform illumination resulting from the optical fiber light delivery, each channel is independently rescaled to have same root-mean-squared (rms) noise, emulating uniform excitation light delivery along the imager. Time-domain results are plotted as summations of groups of 16 adjacent pixels (block of 2×8). For the single-neuron data, to determine the time-domain traces at a voxel source, the processed imaging data at each channel is backpropagated using the ***A^+^*** matrix calculated from the BSS analysis. At the end of the recording session the animals were sacrificed by cervical dislocation.

## ACKNOWLEDGEMENT

This work was supported by the Defense Advanced Research Projects Agency under Contract N66001-17-C-4012 and by the National Science Foundation under Grant 1706207. We gratefully acknowledge TSMC for chip fabrication and their support in the use of experimental SPAD devices.

## AUTHOR CONTRIBUTIONS

S.Y., J.C. and K.L.S. designed the research. J.C. and A.J.T. designed the circuits. S.Y., J.C. and H.Y. performed silicon post-fabrication and optical packaging. A.J.T., J.C. and S.Y. designed the state machine and electronic interface. S.Y. wrote the computational algorithm. J.D.F. helped with device inspection and characterization. A.A., V.A.P., and I.U. designed the animal experiments. V.A.P. performed surgical procedures and prepared eGFP slices. S.Y., I.U. and J.K. performed the *in-vivo* GCaMP and *in-vivo* eGFP mouse experiments. S.Y., J.C., I.U. and J.K. performed data analysis. S.Y., J.C., I.U. and K.L.S. wrote the manuscript and all authors provided comments and edited. L.M., M.L.R. and K.L.S. provided overall supervision and guidance.

## DATA AVAILABILITY

All measurement data relevant to the figures presented in this paper are available at https://github.com/klshepard/acus. All other relevant data are available from the corresponding authors upon reasonable request.

## CODE AVAILABILITY

All scripts used for data analysis are available at https://github.com/klshepard/acus. All other relevant codes are available from the corresponding authors upon reasonable request.

## SUPPORTING INFORMATION

Supporting information includes Figures S1 through S20, Table S1, Sections S1 through S5, Supplementary Movies S1 through S9.

---

**Extended Data Fig. 1.**
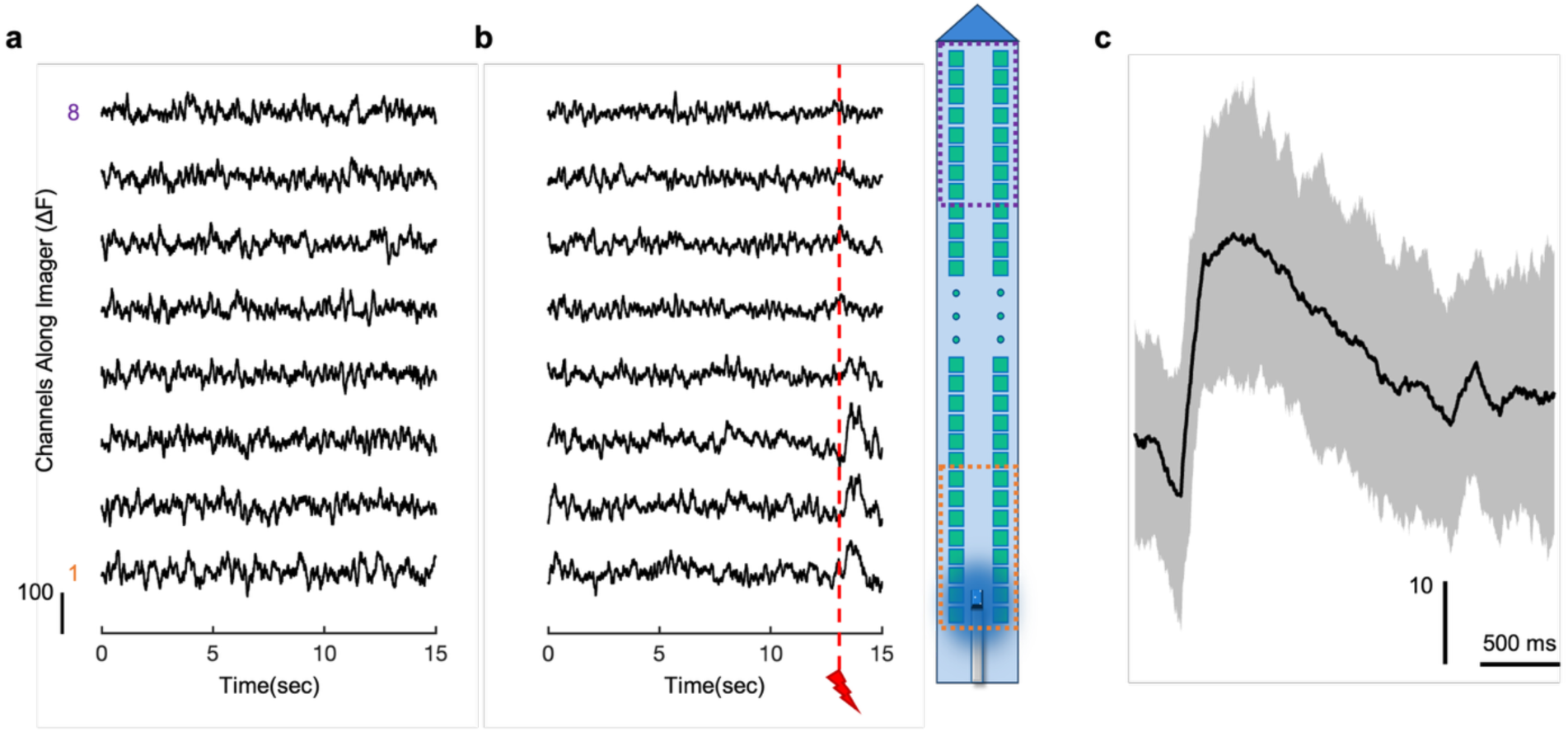
Analysis of *in-vivo* GCaMP6s imaging data. **a,** Photon count acquired by Acus at 400 frames/sec when there is no fluorescent cell nearby. The plot shows the Acus array divided into eight channel groups each of which is a summation of 16 pixels (block of 2×8) as illustrated on the right. **b,** Photon count acquired by Acus at 400 frames/sec when spontaneous activity of an SST cell is detected. The time of electrical micro-stimulation event is noted with the red dashed line. Channels are independently rescaled, detrended and processed with a moving average filter prior to summation for both **a** and **b**, as in Fig. 5e. **c,** Average waveform of calcium activity detected by Acus (n=97) over 33 different SST cells. Black line and the shaded area show the mean and the standard deviation error, respectively.

**Extended Data Fig. 2.**
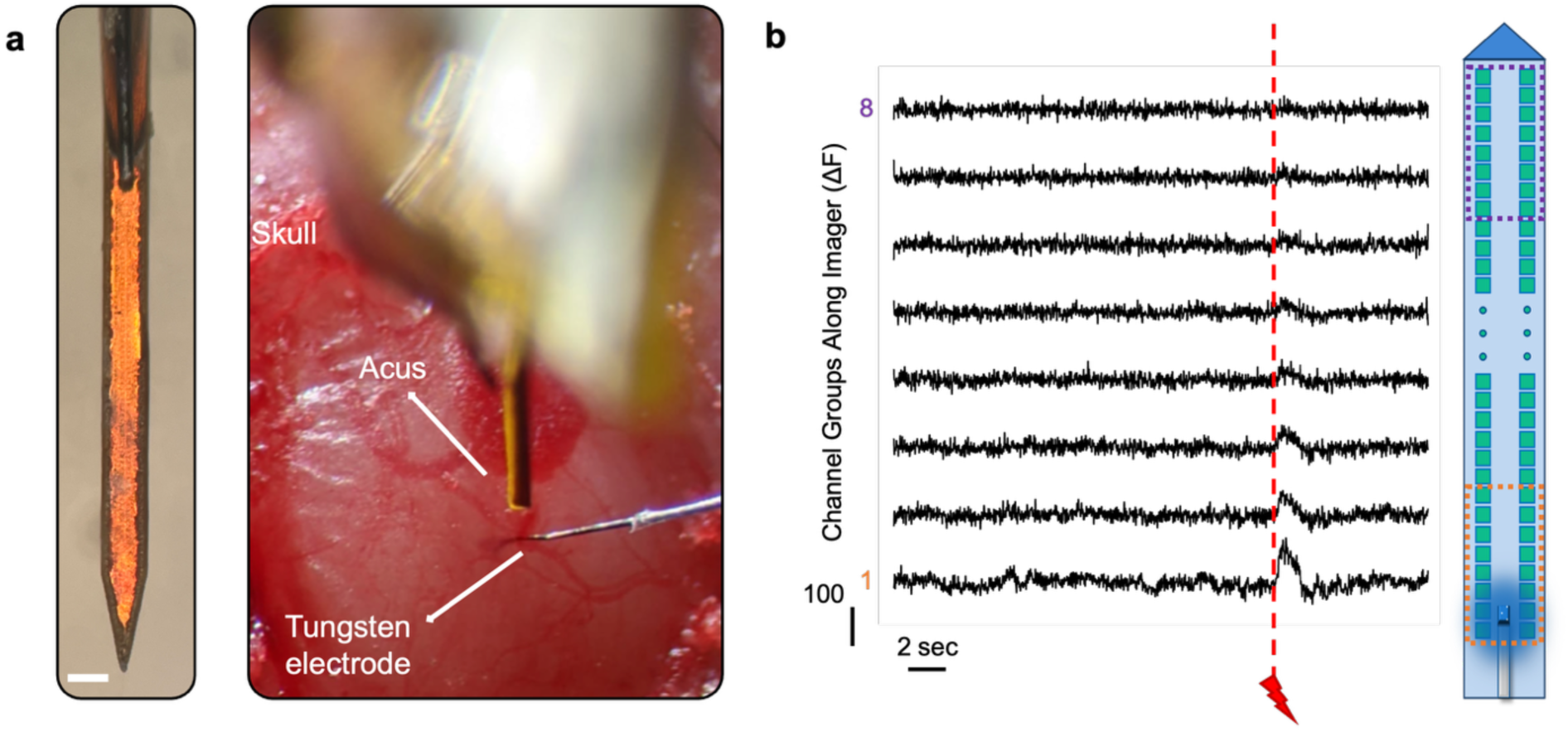
Detection of population activity *in-vivo* with GCaMP6f. **a,** On the left, top view of the fully packaged Acus device, showcasing the shank-based SPAD imager, spectral filters and 50-µm optical fiber (Scale bar: 100 µm). On the right, implantation of Acus and the tungsten electrode in mice cortex. The tungsten electrode is used to deliver 100 µA electrical stimulation. **b,** Photon count acquired by Acus at an integration time of 2.5 ms as a function of time. Each channel group consists of the summation of 16 pixels (block of 2×8) as illustrated on the right. Channels are independently rescaled as in Fig. 5e. The time of the electrical micro-stimulation event is noted with the red dashed line.

