## Supplementary Information for "Implantable CMOS Deep-Brain Fluorescence Imager with Single-Neuron Resolution"

**Supplementary Figures**

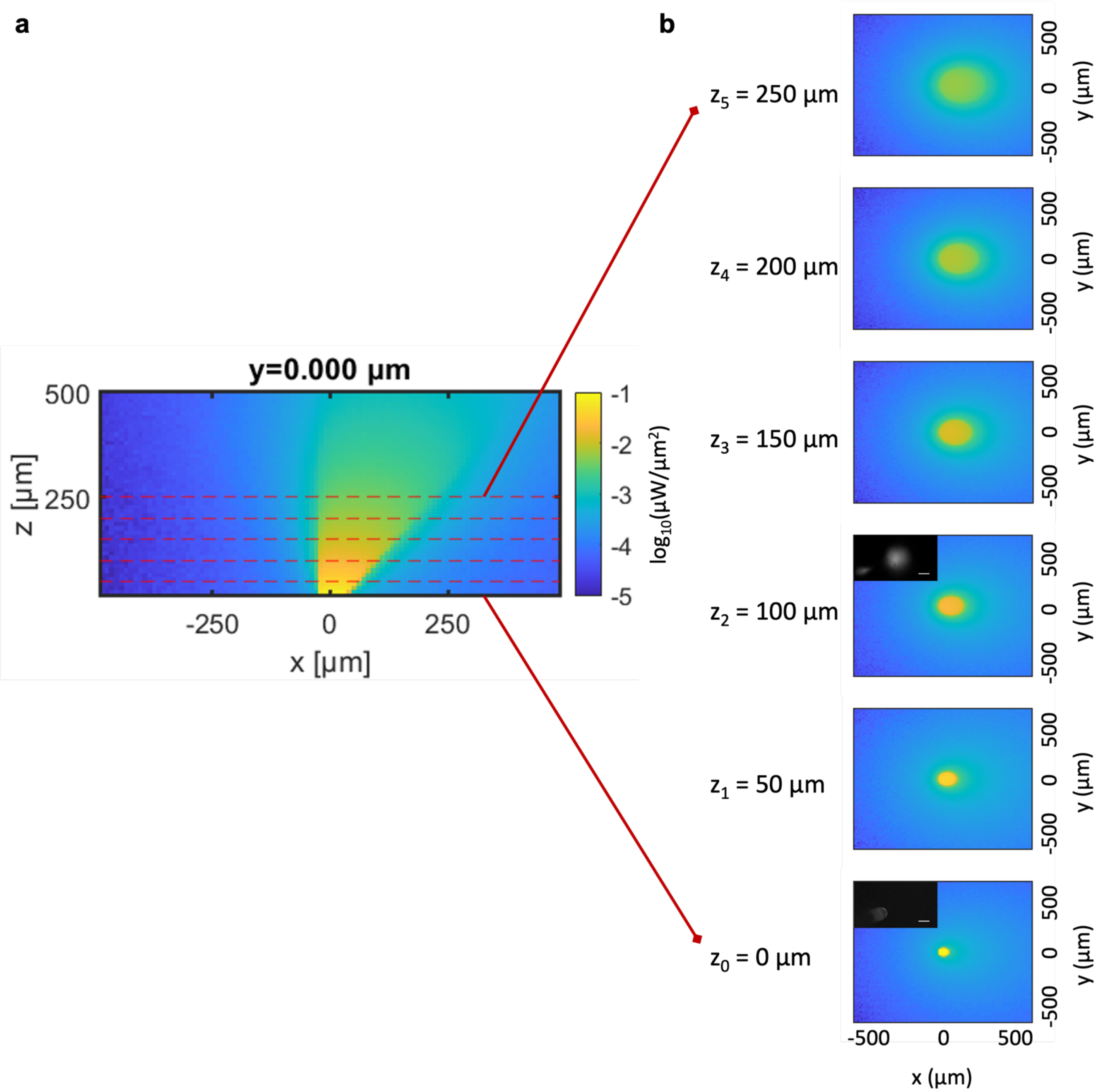

**Supplementary Figure S1. Monte Carlo simulation for angled excitation illumination supplied by Acus during *in vivo* experiments. a,** Monte Carlo photon propagation simulation of 470-nm blue excitation light, supplied by 50-µm optical fiber.[1] Power density at the source is 0.1 µW/µm$^2$. $10^8$ photons were traced in total by using Henyey-Greenstein scattering model with an anisotropy (g) of 0.887, mean scattering length ($L_s$) of 89.9 µm, and mean absorption length ($L_a$) of 16.9 mm, and index of refraction (n) of 1.37. Power density distribution at the center of the y-axis is shown on the left side as xz-graph. Red dashed lines indicate the z slices. **b,** Power density distributions at z slices shown in a. Insets of $z_0$ and $z_2$ show the confocal images. At $z_0$ = 0 µm, power density is 0.1 μW/μm$^2$; at $z_5$ = 250 μm, power density is 3.7 nW/μm$^2$. All plots in a and b share the same colorbar. Scale bars of the inset confocal images represent 50 μm.

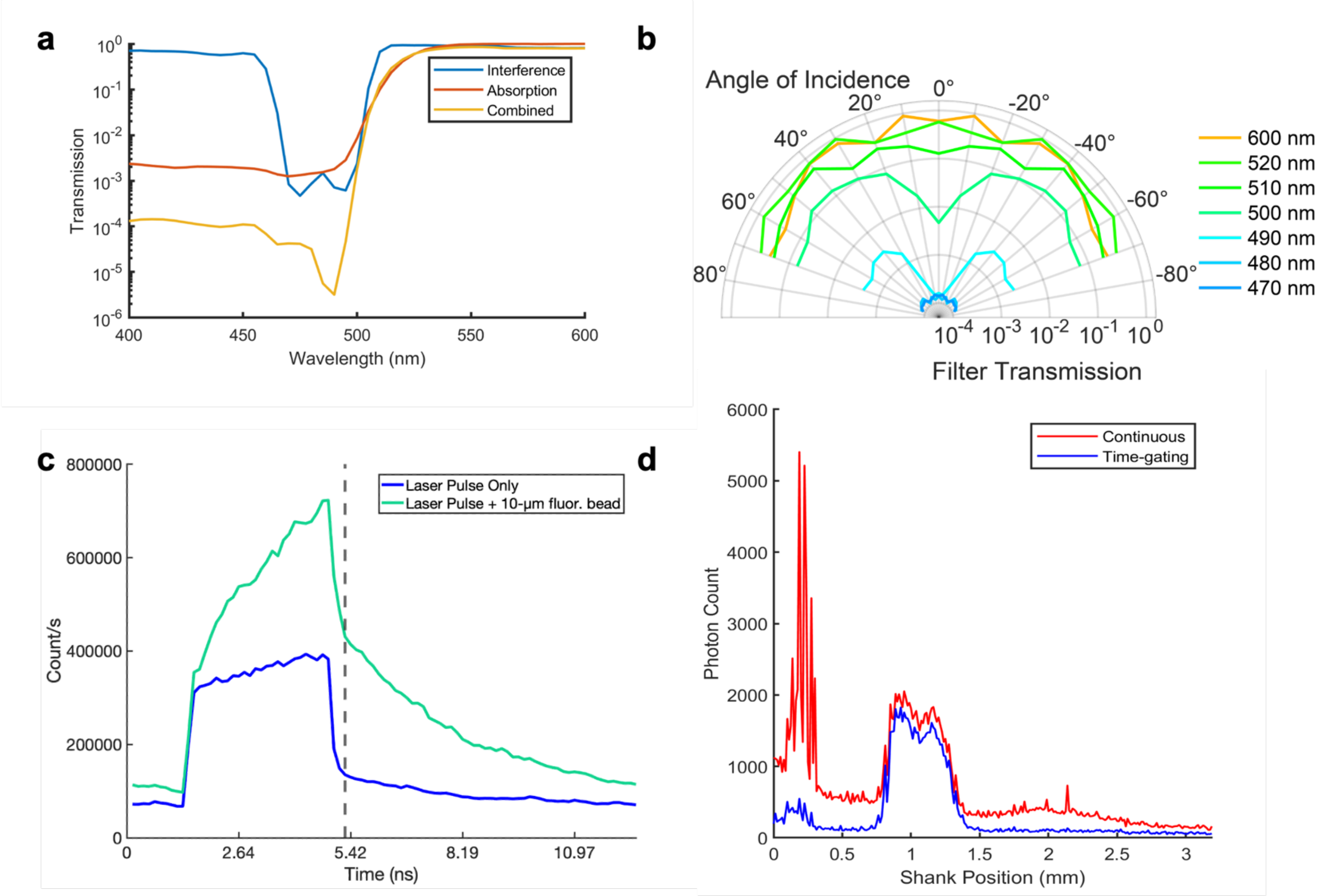

**Supplementary Figure S2. Filter characteristics.**[2] **a,** Interference and absorption filter characterization. **b,** Angular dependency of the Acus pixels after filter deposition as a function of wavelengths. **c,** Impulse response of the SPAD ($t_{rise} = t_{fall} = 300$ ps) created by convolution of laser pulse (80 MHz repetition rate with 140-fs pulse width) and SPAD activation signal with 30% duty cycle. Measured photon counts per second with various gate delays are plotted in two cases: First, only laser pulse is present; second, 10-µm fluorescent bead is added to the system. Dashed line indicates the optimal time-gating position where the excitation pulse is below the 10% of the peak value. **d,** Performance of time-gate filtering resulting in reduced excitation pulse detection.

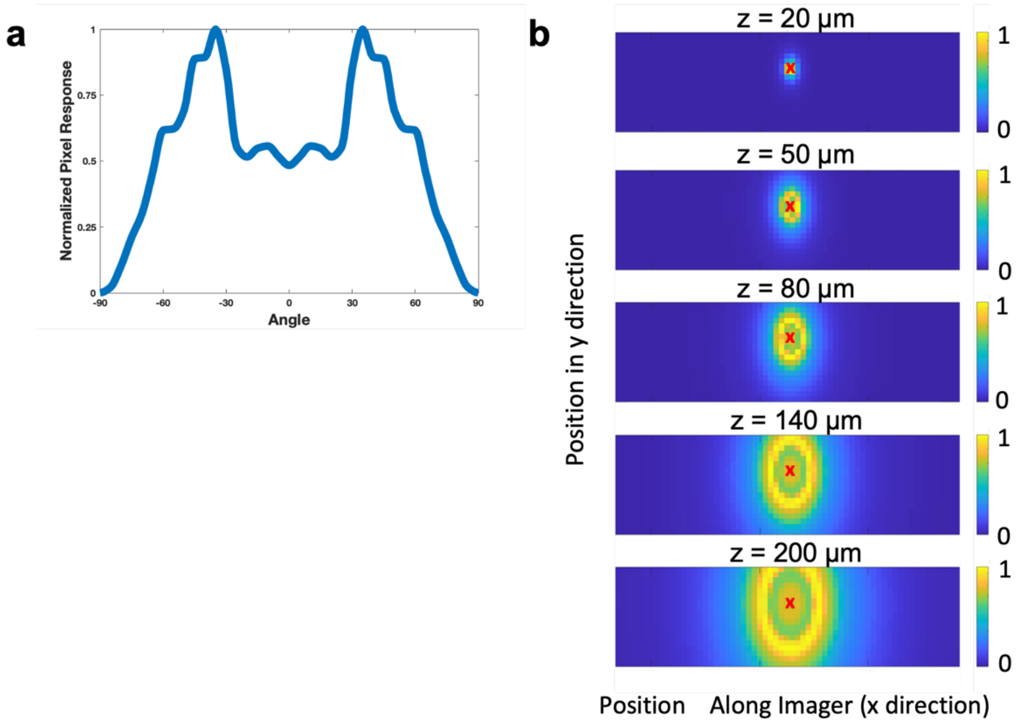

**Supplementary Figure S3. Angular sensitivity of the imager-filter stack. a,** The angular sensitivity profile of SPADs with interference and absorption filter deposited. **b**, Normalized angular sensitivity for a single SPAD pixel (#129) at several imaging planes above the imager array. Pixel location is marked with red "X".

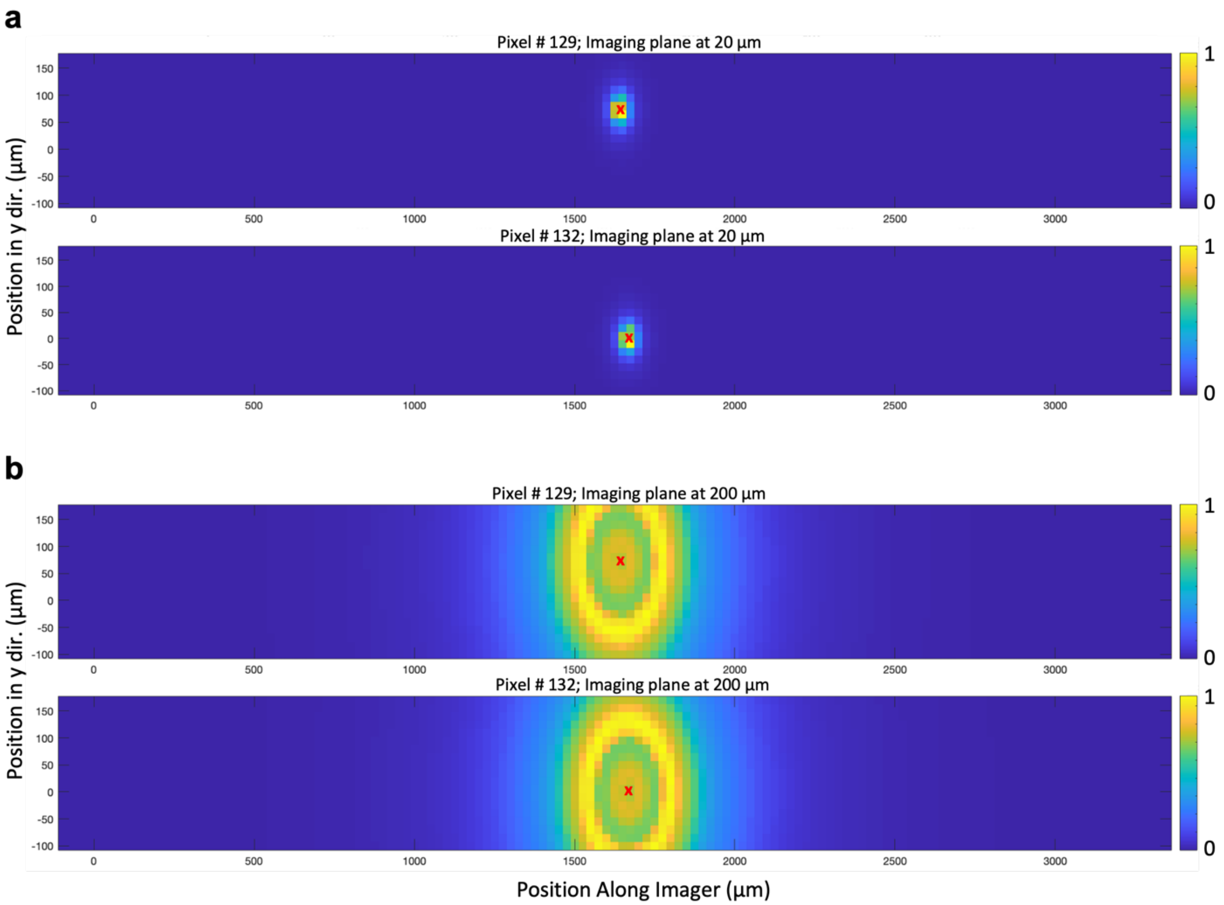

**Supplementary Figure S4. Pixel-to-pixel variability in SPAD angular sensitivity. a,** Angular sensitivity at z=20 µm for adjacent pixels #129 and #132. These pixels are separated by 25.3 µm in *x* and 75 µm in *y*. **b,** Angular modulation at z = 200 µm for pixels #129 and #132. Each pixel location is marked with red "X".

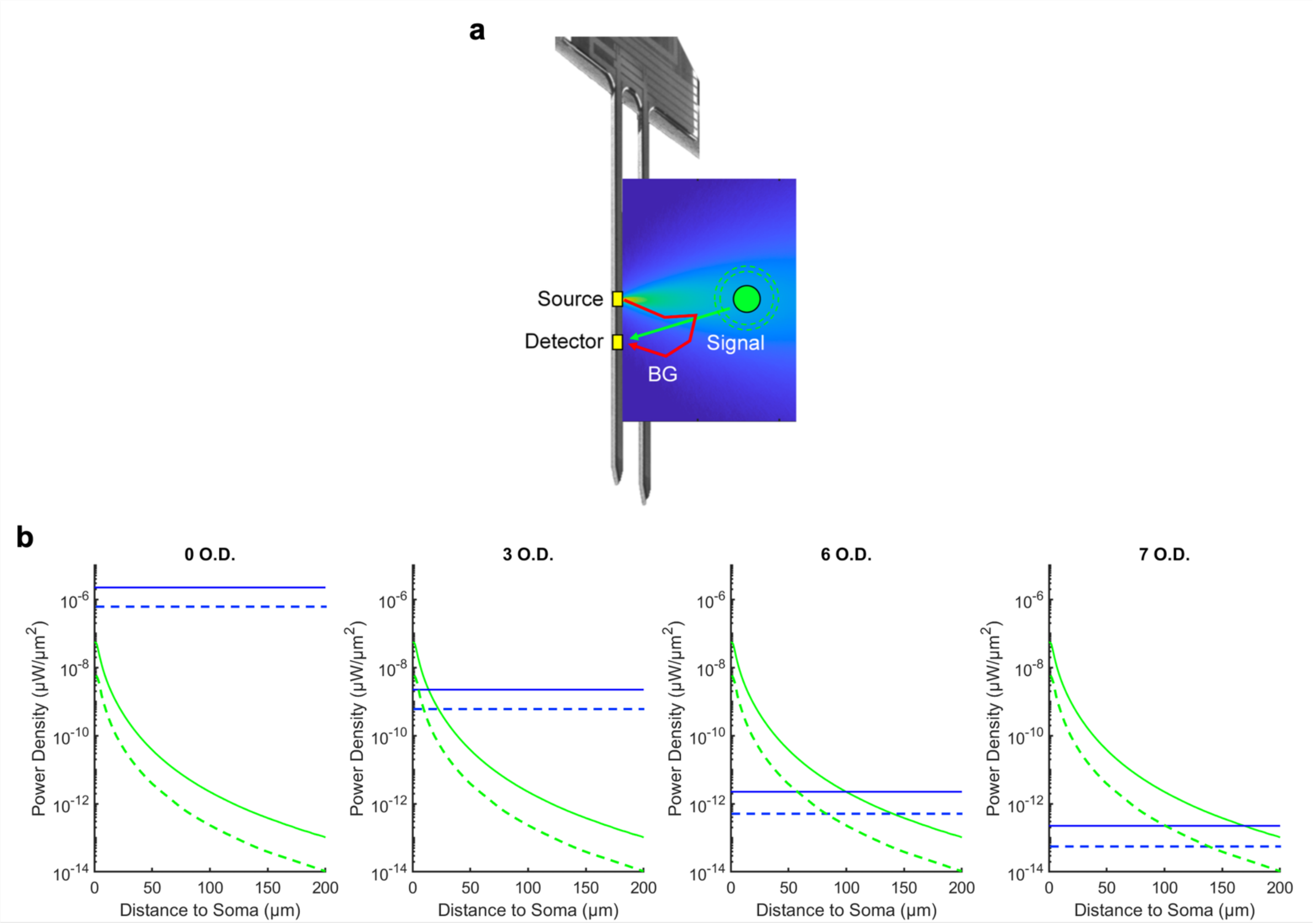

**Supplementary Figure S5. Monte Carlo simulation for filter requirements.**[2] **a,** Monte Carlo photon propagation simulation of a Gaussian-shaped excitation source illuminating neural tissue using the parameters mentioned in Supplementary Figure S1. A total of $10^8$ photons were traced from a 0.1-1 μW/μm$^2$ power sources.[3] The red arrow denotes excitation backscatter, while the green arrow denotes fluorescence emission from a neuronal soma. Detector is assumed to be 25 μm away from the source. **b,** Fluorescence intensity from soma (green) and backscattered excitation intensity from the light source (blue) as a function of increasing distance between soma and the implantable imager. Two power levels are considered: 0.1 μW/μm$^2$ (dashed lines) and 1 μW/μm$^2$ (solid lines). Four different filter O.D. values are compared.

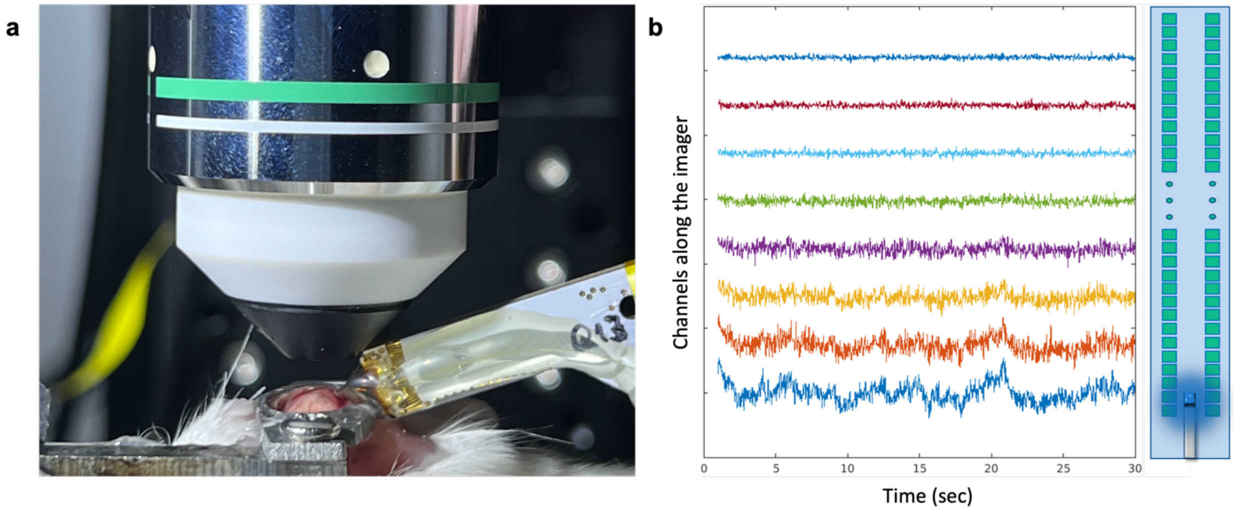

**Supplementary Figure S6.** ***In vivo*** **functional imaging of local field potentials (LFPs) with transgenic GCaMP-expressing mice. a,** Acus insertion into the mouse brain cortex while imaging from the top with a two-photon microscope objective. **b,** Thirty seconds of measured spontaneous LFPs from Acus in the absence of electrical stimulation. Effective channels are formed by integrating 8-by-2 arrays of SPAD pixels.

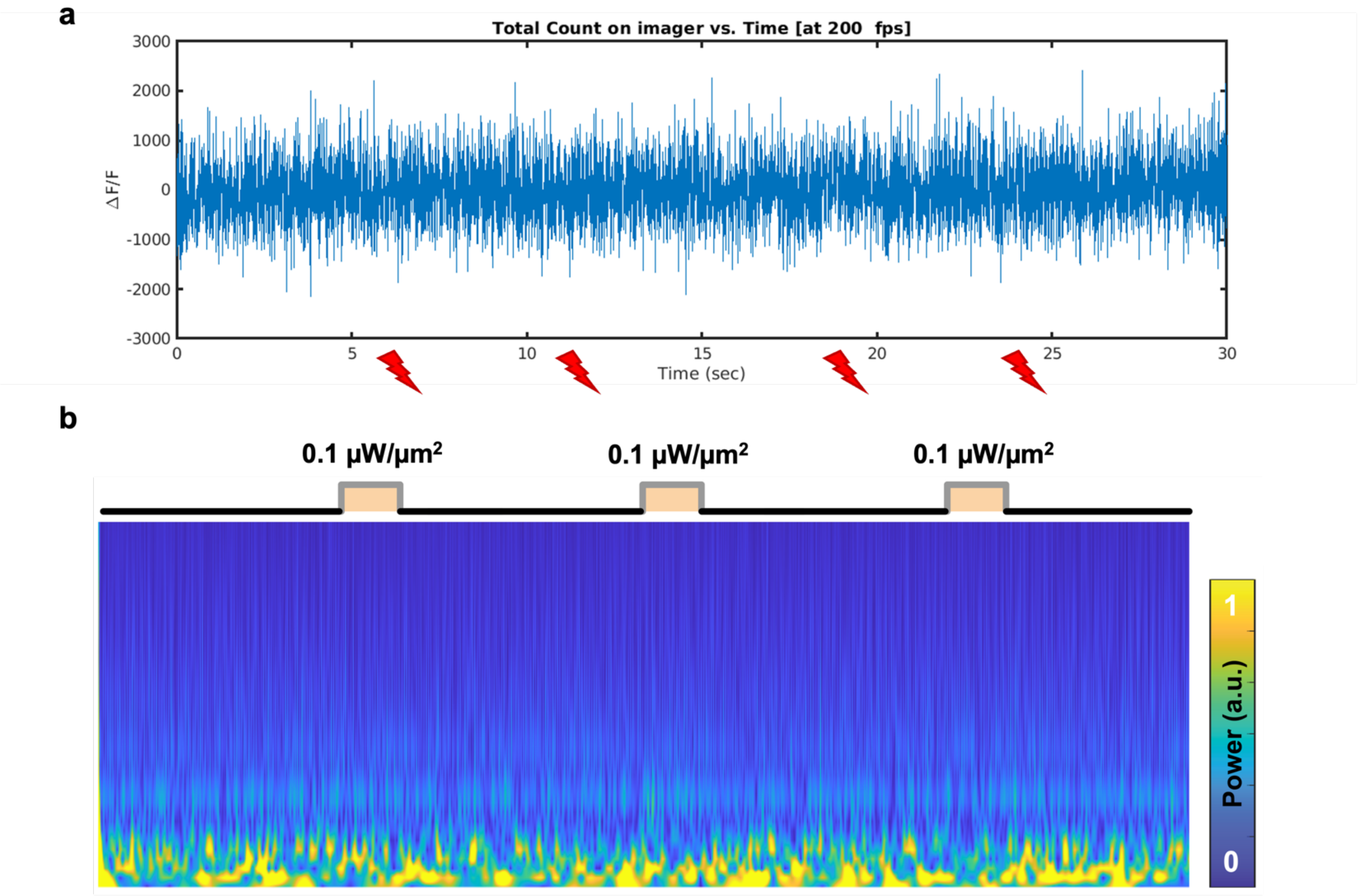

**Supplementary Figure S7. *In vivo* control experiments with wild-type animal. a,** Acus imaging at 200 fps frame rate for thirty seconds during which four 100-µA electrical stimulations, identical to the one in Fig. 5d, are applied with inserted tungsten electrode located 50 µm away from the region of interest in the deep brain. No correlated response is measured by the imager. **b,** Time-frequency graph of electrical recordings from the tungsten electrode implanted in the visual cortex of wild-type mice. The square pulses represent the time of applied excitation light. The bottom colormap illustrates the corresponding LFP power spectra within the 0–50 Hz range, showing no correlated neural response with the laser pulses.

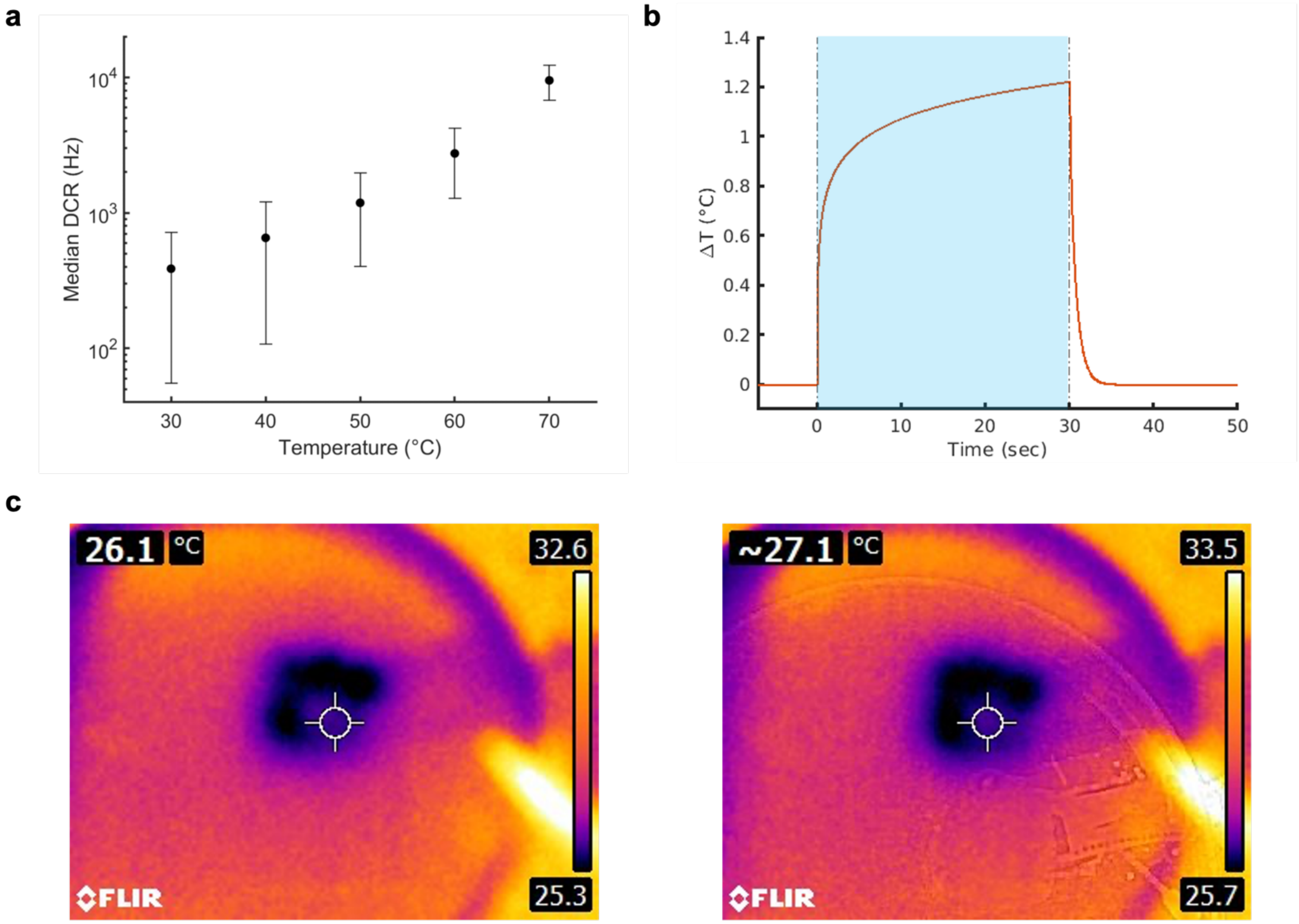

**Supplementary Figure S8. Temperature Characterizations. a,** Dark count rate as a function of temperature over 512 SPAD pixels. Datapoints indicate the median, and error bars indicate the standard deviation.[2] **b,** Response of the logarithmic heat dissipation model to 0.1 μW/μm$^2$ blue excitation light for 30 seconds.[4] **c,** Validation of the heat dissipation model by placing a mouse brain slice on Acus with the fiber light source. Measured temperature change agrees with the simulation model. The slice thickness is 300 μm.

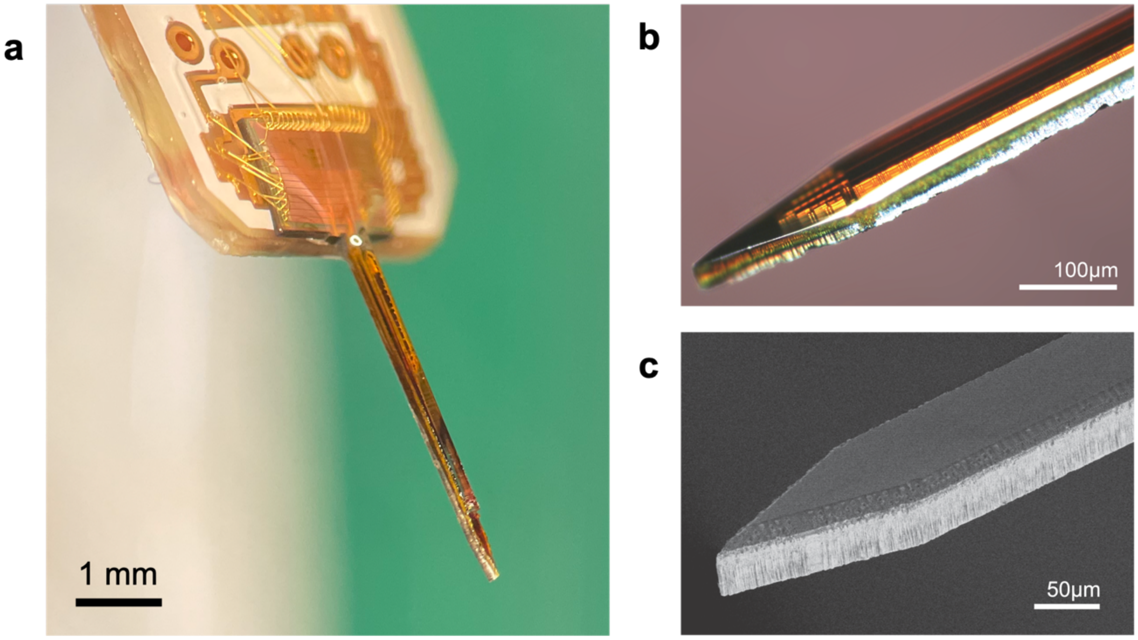

**Supplementary Figure S9. Co-packaging of an optical fiber with the implantable imager.**[2] **a**, Image of the fiber-shank complex prior to insertion. **b,** Magnified image of the distal end of Acus shank at the point of insertion. **c,** Scanning electron microscope (SEM) image of the tip of the Acus shank at the distal end.

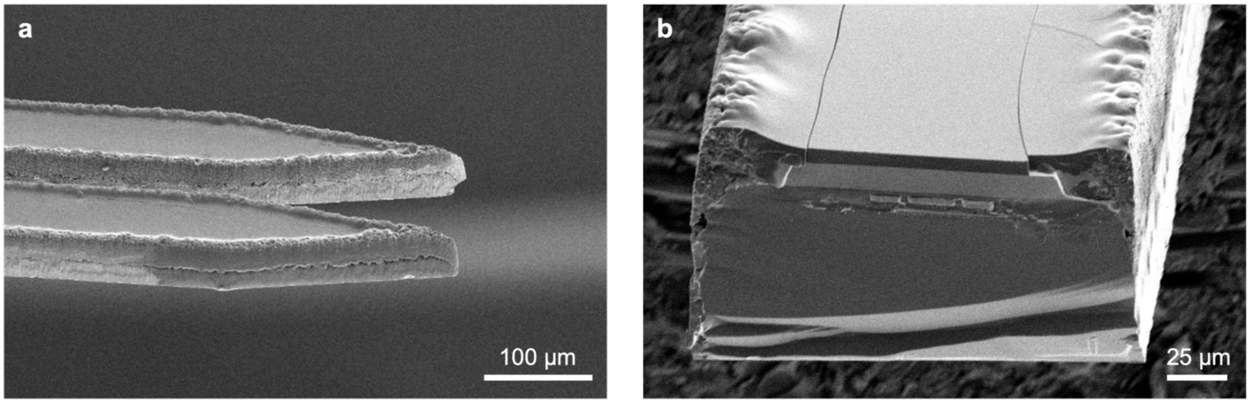

**Supplementary Figure S10. Surface profile created after laser micromachining. a,** SEM profile image of the insertion end of Acus. **b,** SEM section-cut image of the middle of the imager showing the silicon substrate, back-end electronics, and combined spectral filter stack consisting of both the interference filter and absorption filter.

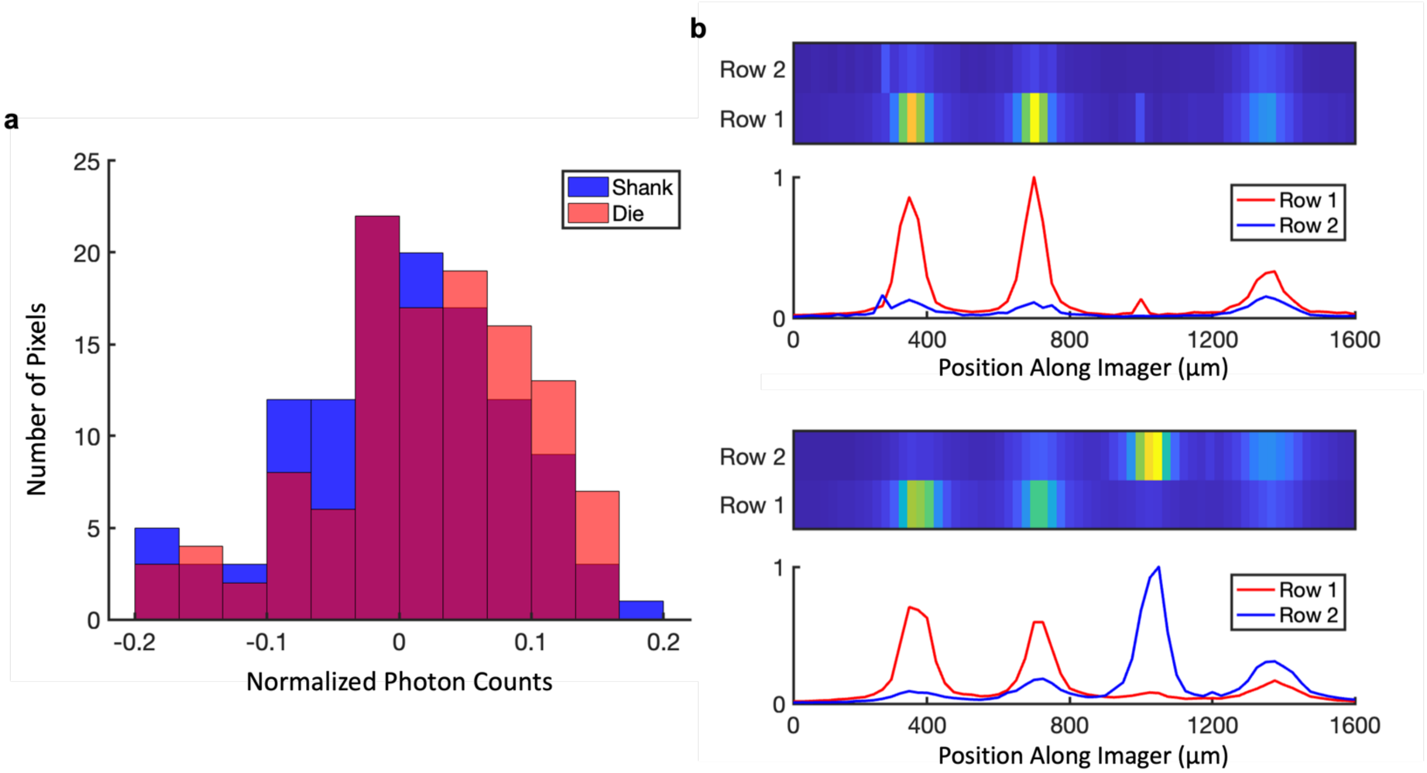

**Supplementary Figure S11. Effect of silicon post-processing on the performance of the imager. a,** Histogram of mean-centered normalized photon counts before ("Die") and after ("Shank") the post-processing, showing that there is no performance degradation due to shank fabrication process. The images used to plot the histograms are taken under identical Gaussian illumination. **b,** Image of three 10-µm fluorescent microspheres taken by the "Shank" (top) and image of four 10-µm fluorescent microspheres taken by the "Die" (bottom) are displayed to show that there is no difference in the measured SNR before and after post-processing.

a

b

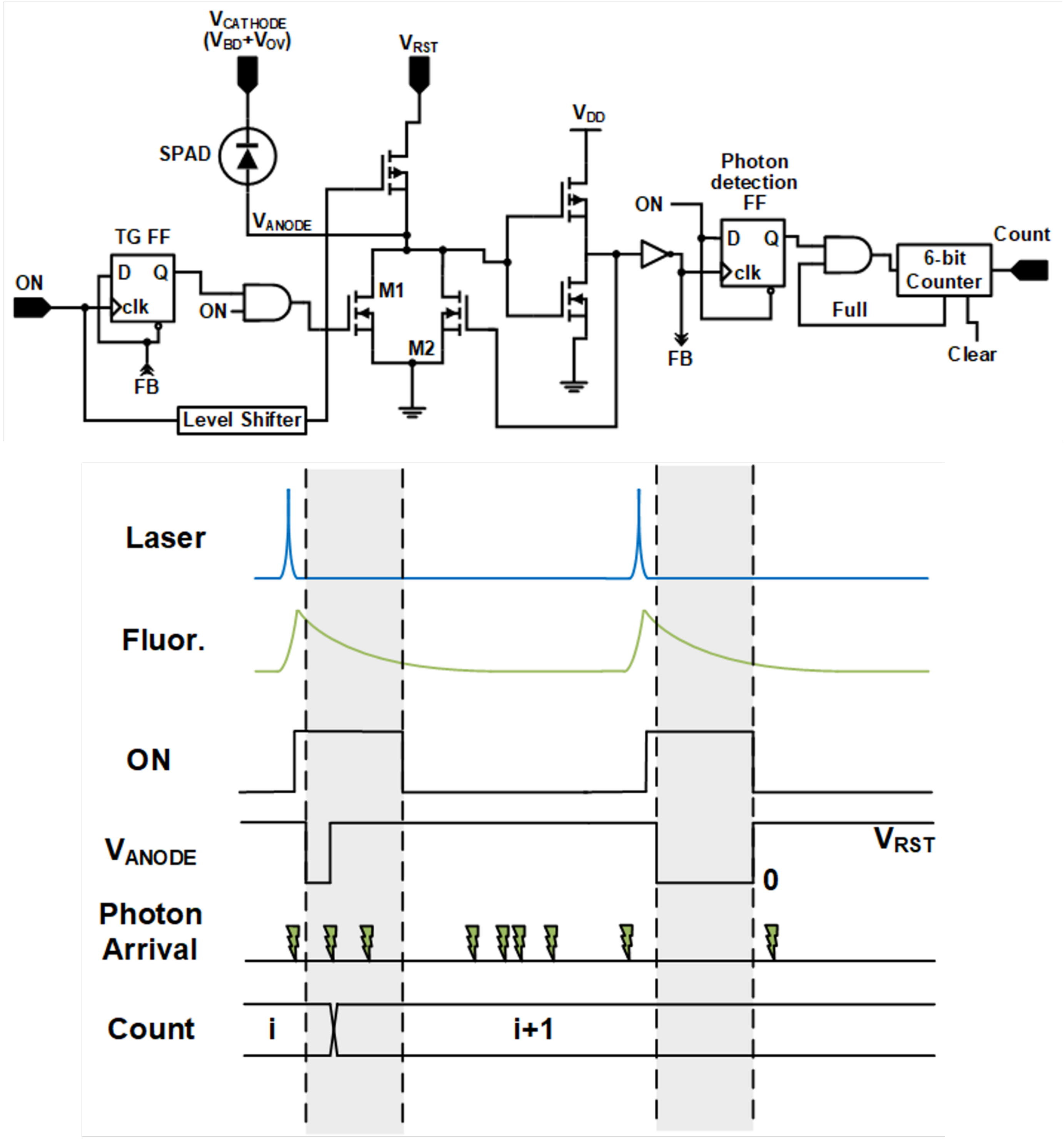

**Supplementary Figure S12. SPAD Pixel Schematics and Operation.**[5] **a,** Per-pixel active-reset quenching circuit with 6-bit counter. **b,** Timing diagram for the time-gated filter in action; the *ON* signal is phase-locked to a pulsed excitation laser, avoiding the collection of excitation light and only collecting fluorescence emission. Effective time-gating window is marked with shaded area where the SPAD is in Geiger mode until the first photon arrival.

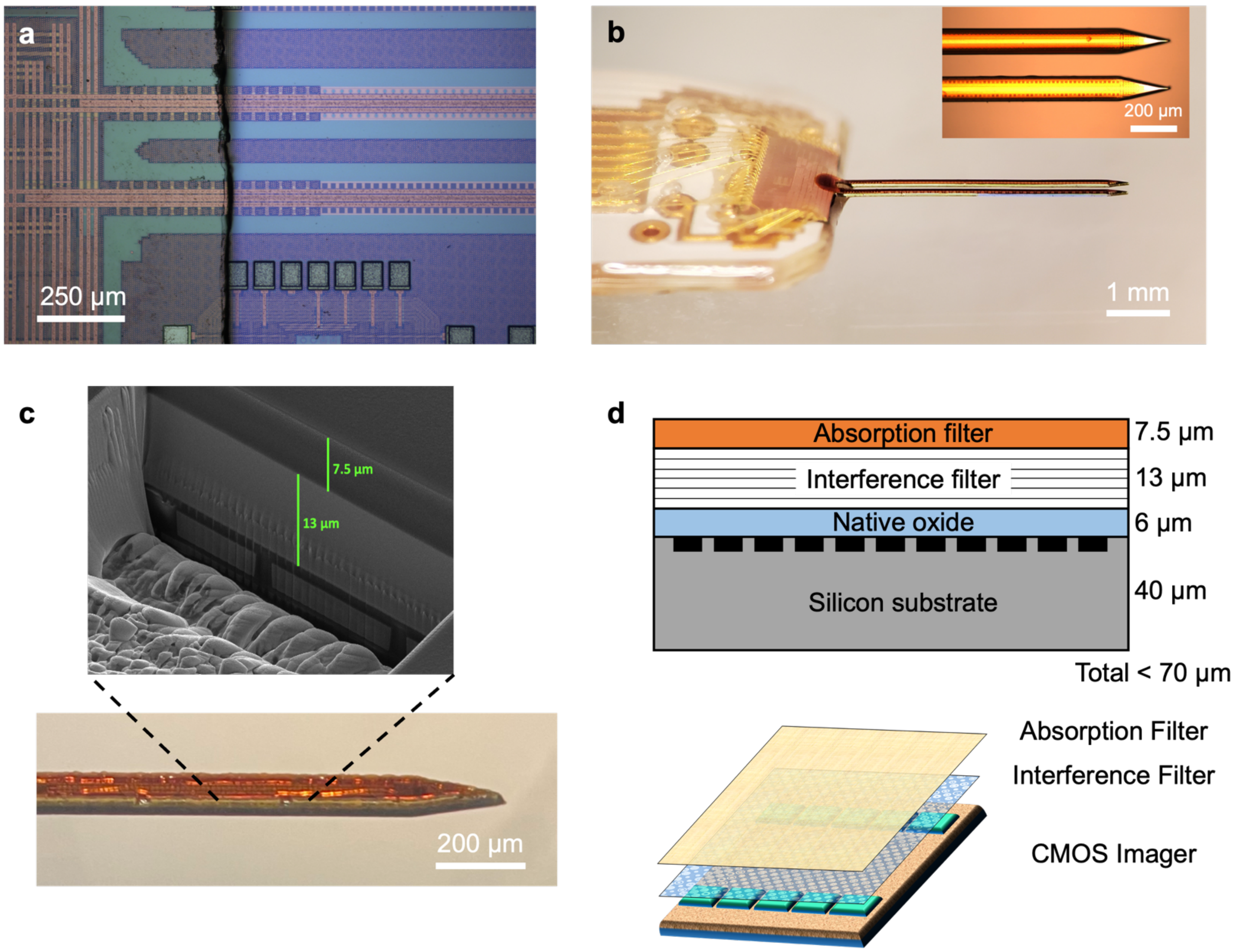

**Supplementary Figure S13. Design of the spectral filters.**[2] **a,** Interference filter deposited on the CMOS chip. **b,** Dye-based absorption filter applied to the shank-format Acus. Inset is showing the magnified image. **c,** Side view of Acus with the combined spectral filter stack. Inset shows SEM image of the spectral filter stack above the back-end metals and their respective thicknesses, 13 μm for the interference filter and 7.5 μm for the absorption filter. **d,** Schematic of the filter stack up with associated thicknesses of each layer.

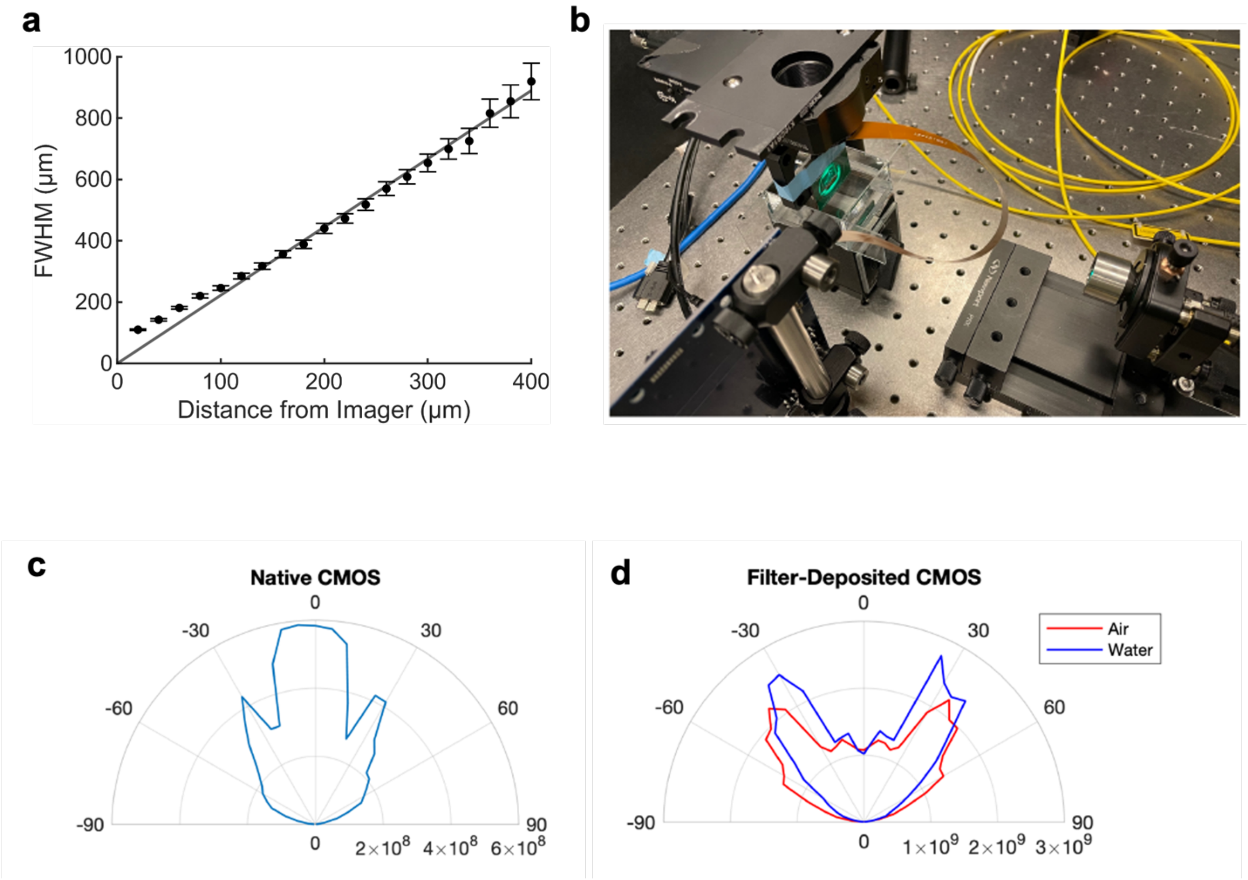

**Supplementary Figure S14. Additional characterization of the SPAD array.**[2] **a,** Full-width-half-maximum change based on the z distance of a single 10-μm fluorescent source. **b,** Angular dependency measurement setup, including collimation of the light and the Acus device on the angle-controlled platform (Thorlabs, Inc.). **c,** Angular modulation of the bare SPAD pixels, out of foundry. **d,** Angular modulation of the SPAD pixels after the deposition of spectral filter stack in two different refractive mediums, air and water.

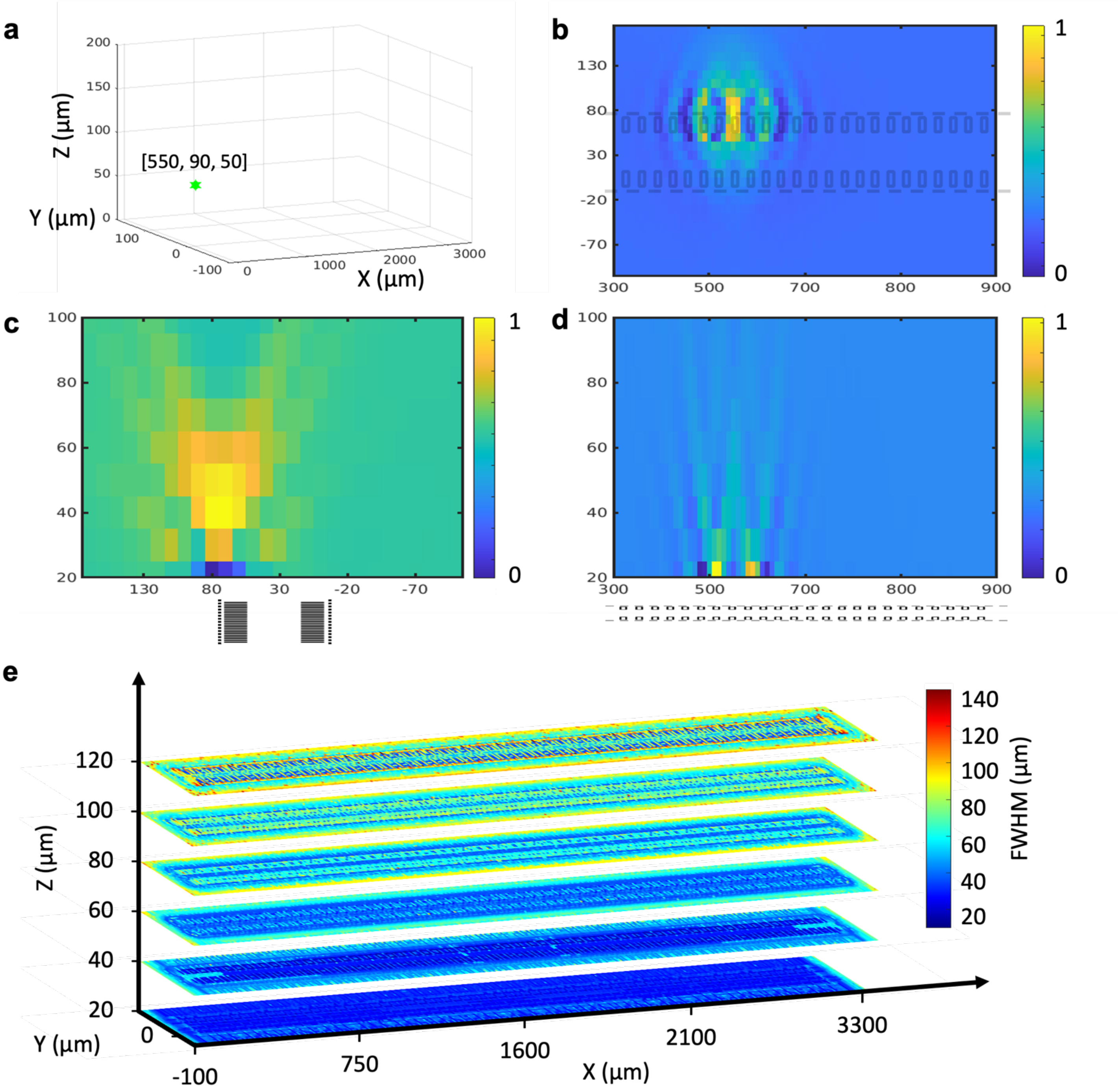

**Supplementary Figure S15. Computational point-spread function (PSF) of the system and resolution at different z distances from Acus. a,** 3D location of the point source used in in the PSF calculation with pseudoinverse backprojection. **b-d,** Computed PSF of an ideal point source located at cartesian coordinates [550, 90, 50 μm], reconstructed with 10-μm voxel resolution. Cross-section of the 3D PSF in x-y, y-z, x-z planes. **e,** FWHM for all voxels calculated by means of Gaussian-fit PSFs in the imaging volume, represented in z-slices. The smallest of PSF FWHM values in the x, y, z directions is chosen as the resolution for the voxel.

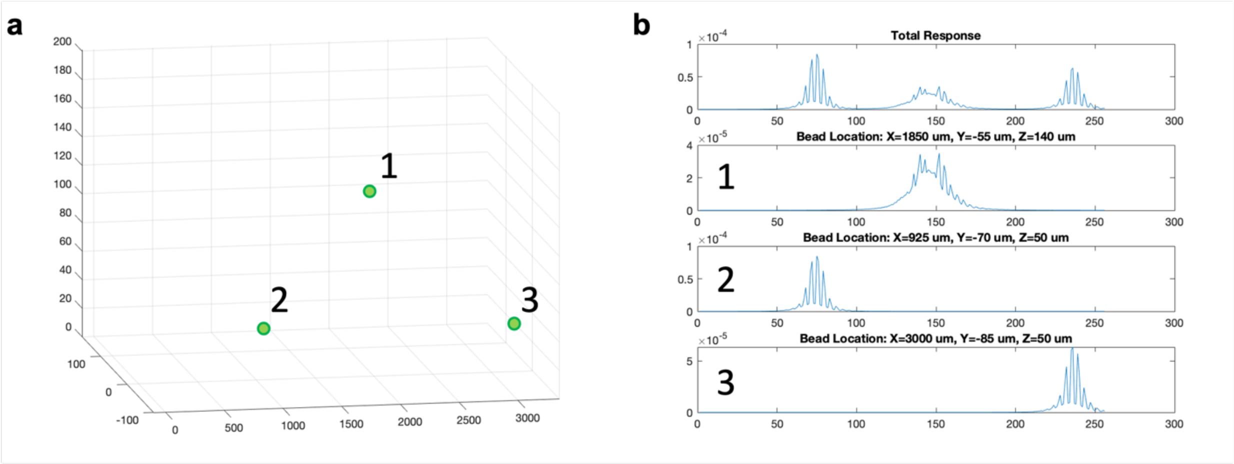

**Supplementary Figure S16. Source generation in 3D and simulated imager response. a,** Three randomly-generated individual fluorescent light sources in the imaging volume of our device. They are labeled as 1, 2 and 3 to demonstrate their individual contributions to the total response in (b). **b,** Expected total linear raw image (256 pixel values are plotted with pixel addressing) at the top and its constituents, labeled as 1, 2 and 3 corresponding to the sources marked in (a).

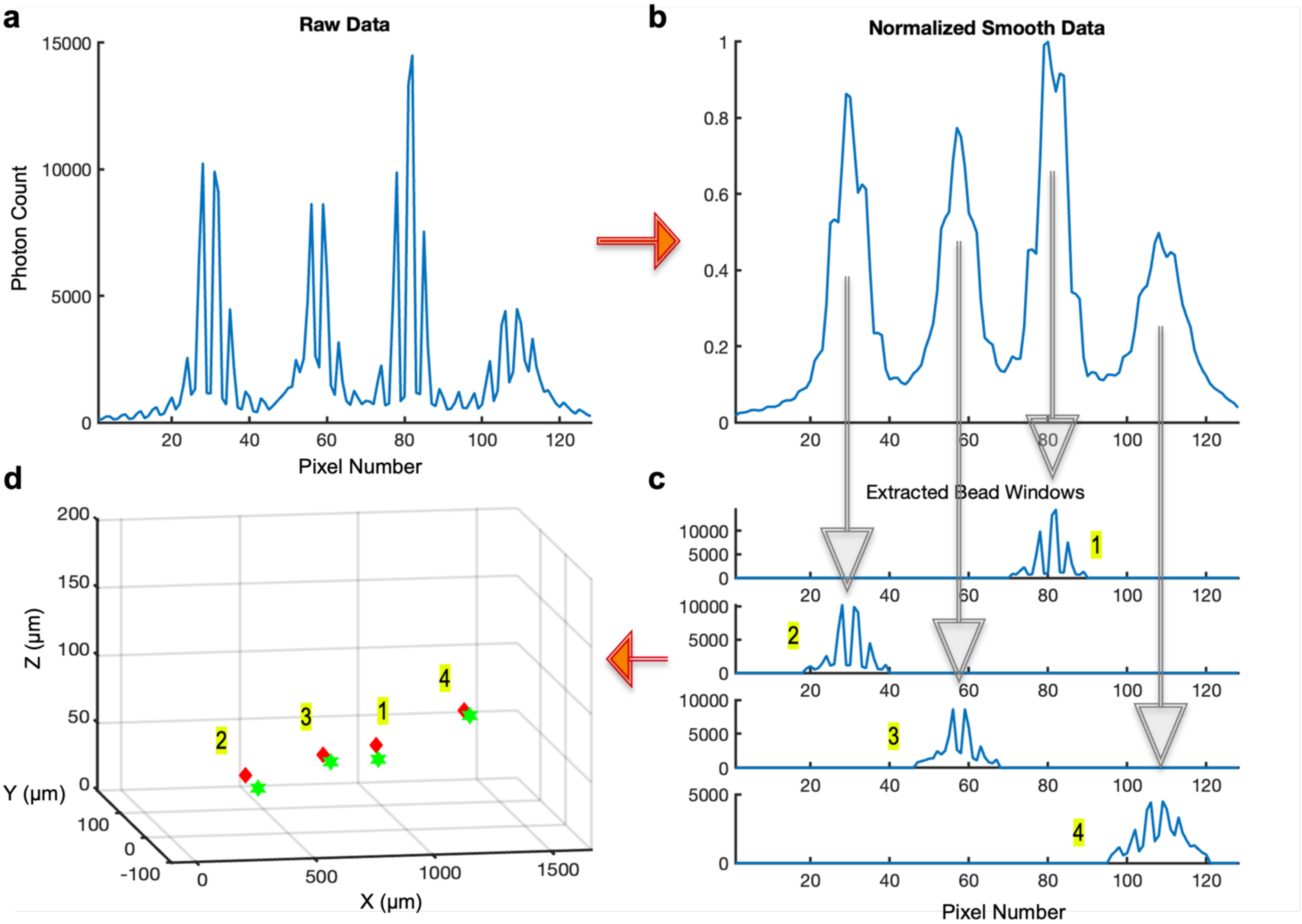

**Supplementary Figure S17. Blind source separation (BSS) algorithm steps on measured data presented in Fig. 3f. a, Raw data generated by 2x64 pixel array when there are four fluorescent sources above the imager**. **b**, To identify regions in the raw image corresponding to the response of different neurons, the raw images are processed with a moving average filter to remove high-spatial-frequency components beyond 50 $\mu m^{-1}$, which is accomplished with a 2×2-pixel moving average, and normalized. Peak detection and width estimation are performed on this "normalized smooth data". Peaks corresponding to separate neurons are detected based on their prominences. For each detected peak, a mask is created based on its location and width. **c**, Separation of individual signals from the raw data using these masks. **d**, Maximum likelihood estimation to determine point source locations. Red and green markers show the true locations and estimated locations of the microspheres, respectively.

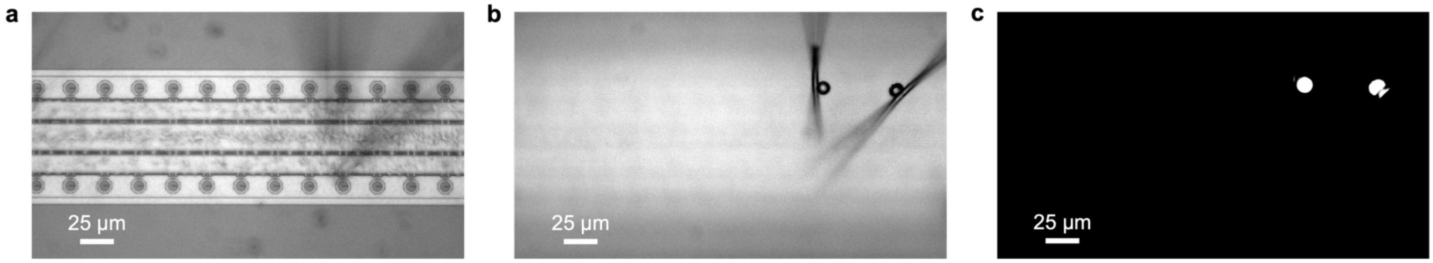

**Supplementary Figure S18. Confocal images from the resolution test conducted with two fluorescent microspheres. a,** Widefield image of the SPAD array consisting of two rows. **b,** Widefield image of the two fluorescent microspheres separated by 50 µm. Custom pipettes are holding the microspheres, hovering 150 µm above the imager. **c,** Fluorescence image of the microspheres shown in b.

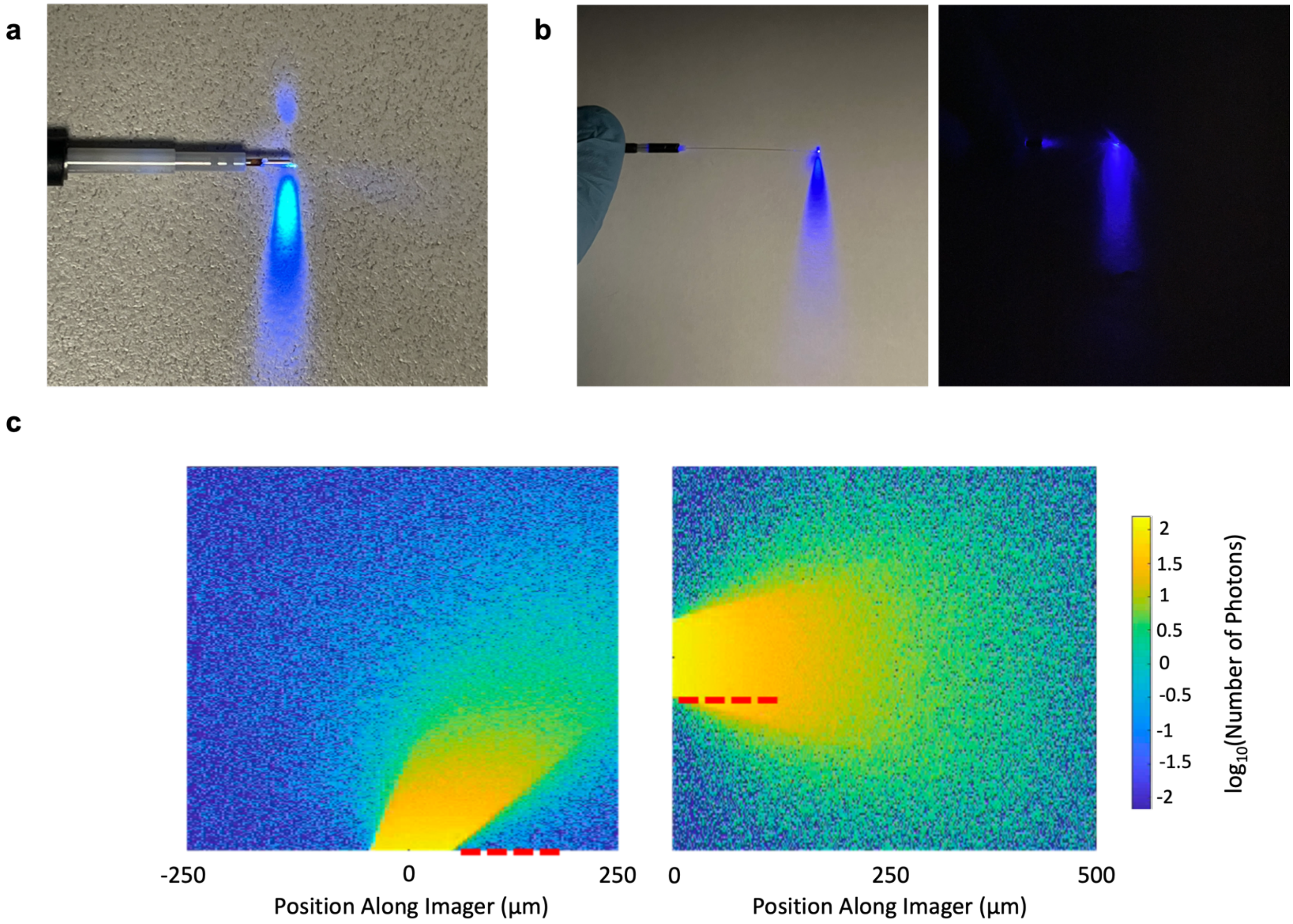

**Supplementary Figure S19. Effect of black epoxy on optical fiber leakage. a,** Before the black epoxy. **b,** After the black epoxy under a light source (left) and in the dark (right). **c,** Monte Carlo simulation for two fiber configurations. On the left, Acus configuration with angled fiber illumination is shown. On the right, configuration with parallel fiber illumination is plotted. Red dashes indicate the four SPAD pixels that are closest to the fibers.

a b

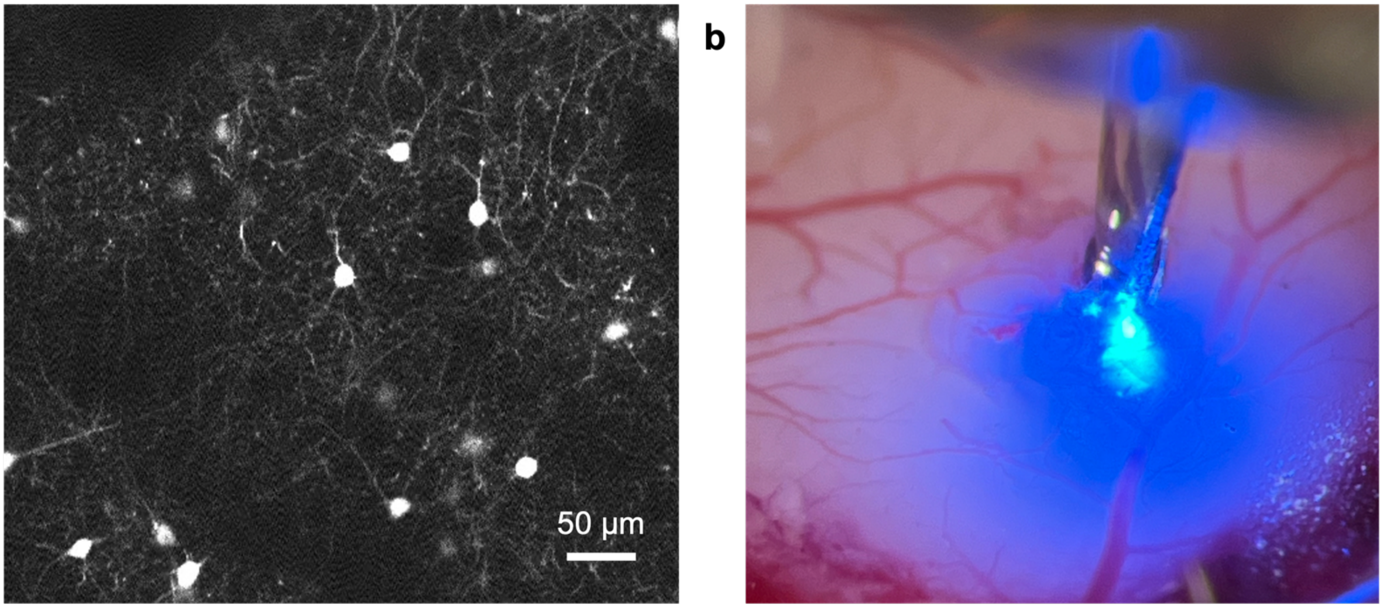

**Supplementary Figure S20.** ***In vivo*** **experiments with eGFP mice. a,** Two-photon images of the GFP-induced mouse brain cortex showing the individual neurons and the sparsity of expression. **b,** Widefield image of the fluorescent excitation of the mouse brain cortex with Acus inserted into the brain.

**Supplementary Table**

| | Field[6] | Perenzoni[7] | Zhang[8] | Ota[9] | Henderson[10] | Acus |
|---|---|---|---|---|---|---|
| **Technology** | 0.13 µm | 0.35 µm | 0.18 µm | 90 nm/40 nm (3D BSI CMOS) | 90nm/40 nm (3D BSI CMOS) | 0.13 µm |
| **Pixel Pitch** | 48 µm | 50 µm | 28.5 µm | 9.6 µm | 9.2 µm | 25.3 µm |
| **Pixel Count** | 64 × 64 | 160 × 120 | 32 × 32 | 960 × 960 | 256 × 256 | 4 × 128 |
| **Power** | 26 W | 157 mW | 310 mW | 370 mW | 77.6 mW | 6.24 mW |
| **Temporal Resolution** | 62.5 ps | 194 ps | 106 ps | - | 35 ps | 137 ps |
| **DCR Median** | 544 Hz @ 1.5 V | 580 Hz @ 3 V | 113 Hz @ 5 V | 2.5 Hz @ 2.5 V | 20 @ 1.5 V | 40 Hz @ 1 V |
| **Max PDP** | 30% @ 1.5V | - | 47.8% @ 5 V | 70% @ 2.5 V | 23% @ 3 V | 12.4% @ 1 V |
| **Fill Factor** | 0.77% | 21% | 28% | ~100% (BSI) | 51% | 6.3% |
| **Frame Rate** | 100 fps | 486 fps | 6 fps | 90 fps | 30-760 fps | 51 kfps |

**Supplementary Table S1. Comparison of implantable fluorescence imager with prior time-gated SPAD imagers.**[5] PDP and DCR are reported at the excess bias voltages indicated; PDP is reported as the maximum value over all wavelengths.

**Supplementary Sections**

### S1. Power dissipation in an implantable imager and heating of the tissue

While power dissipation in traditional microscopes is not a concern, it is of paramount importance in implantable microscopes, which are in direct contact with tissue. In our case, this heating is primarily determined by the light-delivering optical fiber, since the imager on the Acus only consumes an average power of 6.24 mW in typical illumination conditions, resulting in maximum heat flux of 0.8 nW/μm$^2$ across the surface of the device. The use of SPADs is key to this power efficiency since the SPADs themselves only dissipate power when sensing photons (Supplementary Table S1). 6-8 °C change in temperature causes irreversible effect in the brain tissue, and 1.5-3 °C heating results in discernible neural responses.[11,12] We considered both modeling and empirical data to determine heating effects of the Acus.

We used an empirical model to establish a heating simulation for the optical fiber.[4] This model, based on extensive experimental data taken with 500-ms illuminations, has been shown to be more accurate than finite-element models[13,14] and includes the effects of *in-vivo* heat regulation along with the effects of scattering and absorption at different wavelengths at several power densities. With the same power density, smaller fiber outperforms larger ones since there is less chance of hitting a blood vessel, which can become a dominant factor in absorption. At a continuous-wave (CW) power density of 0.1 μW/μm$^2$ from a 50-μm diameter optical fiber at 470-nm wavelength, the temperature graph is given by:

$$f_{rising}(t) = 0.31 \cdot \log(1 + 0.25 \cdot t)$$

where 0.31 is the amplitude constant dominated by the power density and 0.25 is the time constant dominated by fiber tip size. Based on this model, the maximum temperature change will be +0.66°C after 500-ms illumination and +1.22°C after 30-second illumination. The latter is the experimental duration Acus is used for the *in vivo* experiments (Supplementary Fig. S6b). While this model assumes CW illumination, we have a pulse laser system with a pulse frequency of 80 MHz with pulse width of 140 fs, which should result in even less heating. In addition, these models are based entirely on surface measurement, in deep tissue, heating will be reduced due to the ability to conduct heat away omnidirectionally.

Experimentally, we conducted control experiments (Supplementary Fig. S20) to show that there is no heat-induced activity produced by Acus. In addition, we validated our thermal diffusion model by placing a 300-μm-thick mouse brain slice directly on Acus with the attached fiber while monitoring the temperature with a FLIR E5 infrared camera (Teledyne FLIR LLC.). Experimental measurements of the temperature change over time match closely those of the simulations (Supplementary Fig. S6c).

### S2. Deep Reactive Ion Etch (DRIE) of the Acus Shanks

Deep reactive ion etching (DRIE) is used to define the needle-like form factor of Acus from the CMOS die (Supplementary Fig. S7). This trench etching consists of three dry etching steps, a $CF_4$/Ar etch of the thin-film interference filter, a $CF_4$ oxide/nitride etch, and a $SF_6$/$C_4F_8$-based Bosch process silicon etching. Prior to etching, the chip is temporarily bonded with wafer bonding material (Brewer Science, WaferBOND HT-10.11) to a four-inch sapphire wafer. To etch the thin-film interference filter, a CF3/Ar RIE process is used in the Oxford PlasmaPro 100 Cobra (200W RF power/1000W ICP power and 5mTorr pressure with CF4/Ar flow rate, both set as 25sccm). A three-layer photoresist (AZ P4620, Microchemicals GmbH) at a thickness of 30 µm is used as a mask with a 2:1 selectivity for filter-stack over photoresist. Two-layer photoresist (AZ P4620, Microchemicals GmbH) at thickness of 20 µm is used to mask the subsequent oxide/nitride dry-etch process with the same 2:1 selectivity. The oxide/nitride etch is also performed in an Oxford Plasma Pro 100 Cobra (20 W RF power/1000 W ICP power, 10 mTorr pressure with a CF4 flow rate of 45 sccm). The Bosch etch, performed in an Oxford PlasmaLab 80+ ICP, extends approximately 120 µm into the substrate. The trench-etched wafer is fixed on a glass substrate, facing down, with an instant adhesive (Loctite 460, Henkel Corporation). Applying mechanical grinding to the back of the wafer, it is then thinned to a thickness of 40-70 µm with an X-Prep precision milling/polishing system (Allied High Tech Products Inc.).

**S3. Implantable CMOS Imager Design[5]**

Acus is segregated into four voltage domains: 1.5 V for digital core logic, 3.3 V for digital input and output, 5 V for SPAD quenching circuits, and >16 V for SPAD cathode biasing for Geiger-mode operation, which requires a high-voltage foundry process. Acus supports frame rates up to 51 kfps with a 100-MHz reference clock. By summation of frames, higher signal-to-noise ratios (SNRs) can be achieved for lower frame rates.

Acus is able to image 10-µm fluorescent microspheres (Fluoresbrite YG Microspheres 10.0µm, Polysciences Inc.) at 400 frames/sec with SNR in excess of 31 dB. While these microspheres are four times brighter than individual GFP-induced neurons, these frame rates will be required for state-of-the-art high-speed voltage reporters.[15] Acus is an imager comparable in width to silicon electrophysiological probes (< 100 µm) with a length sufficient to cover the extent of the cerebral cortex of a mouse brain (~2 mm). Optimizing the design for this minimally invasive form factor and for imaging performance in low light conditions comes at a cost in dynamic range, field-of-view, and fill factor.

The large aspect ratio of the Acus design brings unique challenges to on-chip interconnect for power and data. For power, the challenge is in reducing IR drops at the furthest points on the shank. For data, RC delays can result in position-dependent data transfer latencies that could contribute fixed pattern noise. We address these challenges by both minimizing power consumption (the shank consumes a maximum total power of 6.24 mW) and delivering the voltages uniformly to each pixel via four 10µm-wide, ultra-thick metal rails running through the shank's centerline, resulting in ~6Ω series resistance at the tip. Decoupling capacitors are distributed all over the imager to reduce delta-I power-supply noise. Controlling the power dissipation of the CMOS circuitry also reduced tissue heating *in vivo* to less than 1.2 °C, important to mitigate the possibility of damage to the tissue.[16]

Acus is in direct contact with neural tissue, which has a refractive index of approximately $n_{brain} = 1.4$. Light entering from brain tissue undergoes less refraction entering the device when compared to air, resulting in a narrower viewing cone for each pixel. At the same time, however, Acus experiences reduced Fresnel reflection in tissue than in air, improving light collection from oblique angles. The resulting angular collection function is as shown in Fig. 3e, also taking into account the masking effects of on-chip metals in the vicinity of each pixel and the reflections within hybrid filter stack.

Acus supports time-gated (TG) fluorescence imaging, which is used for additional background rejection by enabling pixels only after the excitation light has been turned off. The background rejection achieved with TG is limited by the fall time of the excitation light, the SPAD impulse response function (IRF), and delayed diffusive scattering of excitation light. All of these times have to be evaluated relative to the fluorescence lifetime. SPAD IRF is degraded by the generation of photogenerated carriers away from the multiplication region that take time to diffuse, leading to a “tail” in the distribution from those photons that are detected with this delay. This diffusive tail allows photons generated by the excitation light to be detected after the gating window closed. In the same manner, any tail on the turn-off of the excitation light also results in unrejected background. This tail is further enhanced by excitation light scattered through the tissue before being captured in the SPADs. For all these reasons, TG temporal filtering alone is not sufficient, requiring the addition of high-performance optical spectral filters.

### S4. SPAD Imager Performance

The fill factor ($FF$) of the Acus imager is defined by:

$$FF = \frac{A_{SPAD}}{A_{Pixel}}$$

where $A_{SPAD}$ is the cross-sectional active area of the SPAD and $A_{pixel}$ is the cross-sectional area of the pixel. The dark current rate (DCR) is defined as observed avalanche rate in the absence of light and afterpulsing. Afterpulsing refers to false events that are correlated in time with previous detection event and are the result of the action of traps which capture photogenerated carriers and subsequently release them. The afterpulsing probability (APP) is determined from the inter-avalanche histogram method.[17] When Acus is used *in vivo*, it operates at an elevated 38°C, which increases the DCR by about 45% compared to room temperature operation. (Supplementary Fig. S6a)

Photon detection probability is defined as the probability of triggering an avalanche in the event of photon absorption in the pixel and is given by:

$$PDP = \frac{Detected\ Photon\ Count - DCR * t_{int}}{Incident\ Photon\ Count} \cdot (1 - APP)$$

where $t_{int}$ is the integration time. Photon detection efficiency ($PDE$) is given by ($PDP$ is the photon detection probability and $FF$ is the fill factor):

$$PDE = PDP \cdot FF$$

After subtraction of the background image, i.e. calculating ΔF/F, there are two main sources of noise that limit the SNR. The first is the photon-shot noise created by the collected fluorescence signal ($P_{sig}$) and the second one is the photon shot noise of the unrejected background excitation light ($B$). Hence the SNR can be calculated as follows:

$$SNR = \frac{P_{sig}}{\sqrt{P_{sig} + B}}$$

where both $P_{sig}$ and $B$ are calculated in detected photoelectrons.

Noise equivalent power (NEP) is the minimum signal intensity required for an SNR of 1 within a 1 Hz bandwidth and is given by ($h$ is Planck's constant, $v$ is the frequency of the photon, $DCR$ is dark count rate, $PDE$ is the photon detection efficiency):

$$NEP = hv \cdot \frac{\sqrt{2 \cdot DCR}}{PDE}$$

Dynamic range as determined by the number of bits (Q) of the counter is given by:

$$DR = 20 \times \log_{10} 2^{Q}$$

### S5. Minimum Filter Requirement

We performed Monte Carlo ray-tracing simulations to determine the degree of excitation rejection required for an embedded optical source and detector in tissue, taking full account that excitation light must backscatter a full 180° in order to be collected which helps to improve background rejection. A 1-µW, 470-nm Gaussian light source (8° half-angle) illuminates a neural tissue model (Supplementary Fig. S5a). The tissue has an 89.9-µm mean scattering length, 16.9-µm absorption length, 1.37 refractive index, and Henyey-Greenstein phase function (anisotropy = 0.887). Fluorescence comes for a 15-µm-diameter neuronal soma containing 10 µM GFP (55,000 $M^{-1}\cdot cm^{-1}$ extinction coefficient, 0.79 quantum yield). In fact, this is an underestimation of the actual fluorescence likely to be observed with GFP labelling, where concentrations may actually be closer to 100 µM.[18] The detector is assumed to be 25 µm from the source on the same plane.

Supplementary Fig. S5b shows the simulated photon fluxes for varying distances between the imager and the modeled neural soma. Fluorescence decreases with increasing distance with scattering and absorption of both the excitation light and fluorescence. Backscattered excitation intensity remains constant regardless of soma position. Four emission filters are compared for their effect on SNR. The photon-shot noise from unrejected background degrades noise performance in the presence of inadequate filtering. A filter with optical density (OD) 3 distinguishes somas within 20 µm, but SNR remains under 100. OD 6 allows the identification of cells at 100-µm distance; improving it to OD 7 extends detectable distance to 170 µm. Each incremental OD gain provides greater readable depth. OD 7 performance is delivered by the hybrid emission filter and time-gating used in Acus.
